# Facilitation of Ca_V_3.2 channel gating in pain pathways reveals a novel mechanism of serum-induced hyperalgesia

**DOI:** 10.1101/2025.01.03.631165

**Authors:** Karolina Sanner, Sarah Kawell, J. Grayson Evans, Vida Elekovic, MacKenzie Walz, Sonja Lj. Joksimovic, Srdjan M. Joksimovic, Rebecca R. Donald, Maja Tomic, Peihan Orestes, Simon Feseha, Annemarie Dedek, Seyed Mohammadreza Ghodsi, Isabella P. Fallon, Jeonghan Lee, Sung Mi Hwang, Sung Jun Hong, John P. Mayer, Douglas F. Covey, Carmelo Romano, Tamara Timic Stamenic, Jean Chemin, Emmanuel Bourinet, Gaetan Poulen, Nicolas Longon, Florence Vachiery-Lahaye, Lue Bauchet, Charles F. Zorumski, Michael H. B. Stowell, Michael E. Hildebrand, Elan Z. Eisenmesser, Vesna Jevtovic-Todorovic, Slobodan M. Todorovic

## Abstract

The Ca_V_3.2 isoform of T-type voltage-gated calcium channels plays a crucial role in regulating the excitability of nociceptive neurons; the endogenous molecules that modulate its activity, however, remain poorly understood. Here, we used serum proteomics and patch-clamp physiology to discover a novel peptide albumin (1-26) that facilitates channel gating by chelating trace metals that tonically inhibit Ca_V_3.2 *via* H191 residue. Importantly, serum also potently modulated T-currents in human and rodent dorsal root ganglion (DRG) neurons. *In vivo* pain studies revealed that injections of serum and albumin (1-26) peptide resulted in robust mechanical and heat hypersensitivity. This hypersensitivity was abolished with a T-channel inhibitor, in Ca_V_3.2 null mice and in Ca_V_3.2 H191Q knock-in mice. The discovery of endogenous chelators of trace metals in the serum deepens our understanding of the role of Ca_V_3.2 channels in neuronal hyperexcitability and may facilitate the design of novel analgesics with unique mechanisms of action.

## Introduction

Intractable pain is estimated to affect about 10% of the worldwide population ^1–3^. Unfortunately, current treatments, such as opioids and antiepileptics, are inconsistently effective and are often associated with serious side effects ^2–6^. A deeper understanding of the molecular mechanisms of pain may elucidate alternative targets for the development of novel treatments with more favorable efficacy and safety profiles.

We previously demonstrated that the Ca_V_3.2 subtype of T-type voltage-gated calcium channels (T-channels) profoundly influences the excitability of nociceptive sensory neurons within the dorsal root ganglia (DRG) ^7,8^ and supports glutamate-mediated excitatory synaptic transmission in dorsal horn (DH) neurons ^9^. When activated, Ca_V_3.2 channels produce a depolarizing current at subthreshold membrane potentials, which facilitates the activation of voltage-gated sodium channels (I_Na+_), triggering action potentials and/or glutamate release at presynaptic terminals. Consequently, Ca_V_3.2 channel activation can enhance the likelihood that a nociceptive signal will be transmitted from the periphery to the central nervous system ^10^. In addition to these physiological attributes, Ca_V_3.2 channels are involved in neuronal hyperexcitability in animal models of intractable pain, such as diabetes ^8,11^, peripheral nerve injury ^12^, somatic ^13^ and visceral inflammation ^14^, chemotherapy ^15^, and surgical incision inflammation ^16^. Although these studies establish the importance of post-translational modification in regulating Ca_V_3.2 activity (e.g., glycosylation, de-ubiquitination), the role of endogenous modulators as a cause of T-channel hyperactivity remains unclear.

The DRG are exposed to whole blood, including its components, such as serum, in various physiological and pathological conditions in both humans ^17^ and animals ^18^. The dense vascularization and high permeability of capillaries in the DRG may make sensory neurons susceptible to various toxic, inflammatory, and endogenous mediators, resulting in painful neuropathies ^17^. Here, we used serum proteomic analyses to identify endogenous factors and investigated their effects on both recombinant and native Ca_V_3.2 currents using patch-clamp techniques *in vitro* and *ex vivo*, as well as *in vivo* pain assays. Unexpectedly, we discovered that in addition to serum itself, the N-terminal albumin fragment albumin (1-26) facilitates T-channel gating, increases nociceptor excitability, and induces hyperalgesia *in vivo* by interacting with a metal-binding H191 Cav3.2 residue.

## Results

### Serum increases current densities and facilitates gating of recombinant and native DRG Ca_V_3.2 channels

We investigated the effect of fetal bovine serum (FBS) on recombinant human Ca_V_3.2 current density using patch-clamp recordings. Perfusion of human embryonic kidney (HEK)-293 cells with 1% FBS resulted in a large increase in Ca_V_3.2 current amplitudes over a wide range of test potentials, while washout of FBS reestablished the baseline levels of the currents (Figure 1A-C). Notably, 1% FBS induced a hyperpolarizing shift of approximately 5 mV in the half-maximal voltage-dependence of activation (V_50_) of Ca_V_3.2 currents (Figure 1D). Similarly, 1% FBS shifted the mean V_50_ for steady-state activation by approximately -9 mV in acutely dissociated rat DRG cells (mean V_50_ of controls: -47.8 ± 4.4 mV; mean V_50_ with FBS: -56.8 ± 5.6 mV, n=5, p<0.01, two-tailed paired t-test). In contrast, we found that 1% FBS had little effect on aspects of the voltage-dependent kinetics of Ca_V_3.2 channel inactivation, including steady-state inactivation and recovery from inactivation (data not shown). We also noticed a prominent hyperpolarizing shift in steady-state activation curves and minimal effect on voltage-dependent inactivation process in our experiments with 1% human serum in acutely dissociated rat DRG cells (Supplementary Figure 1). Hence, we concluded that serum preferentially facilitates T-channel gating. Importantly, we found that 1% human serum also robustly increased the amplitude of T-currents in human DRG cells *in vitro* (Supplementary Figure 2).

**Figure 1.**
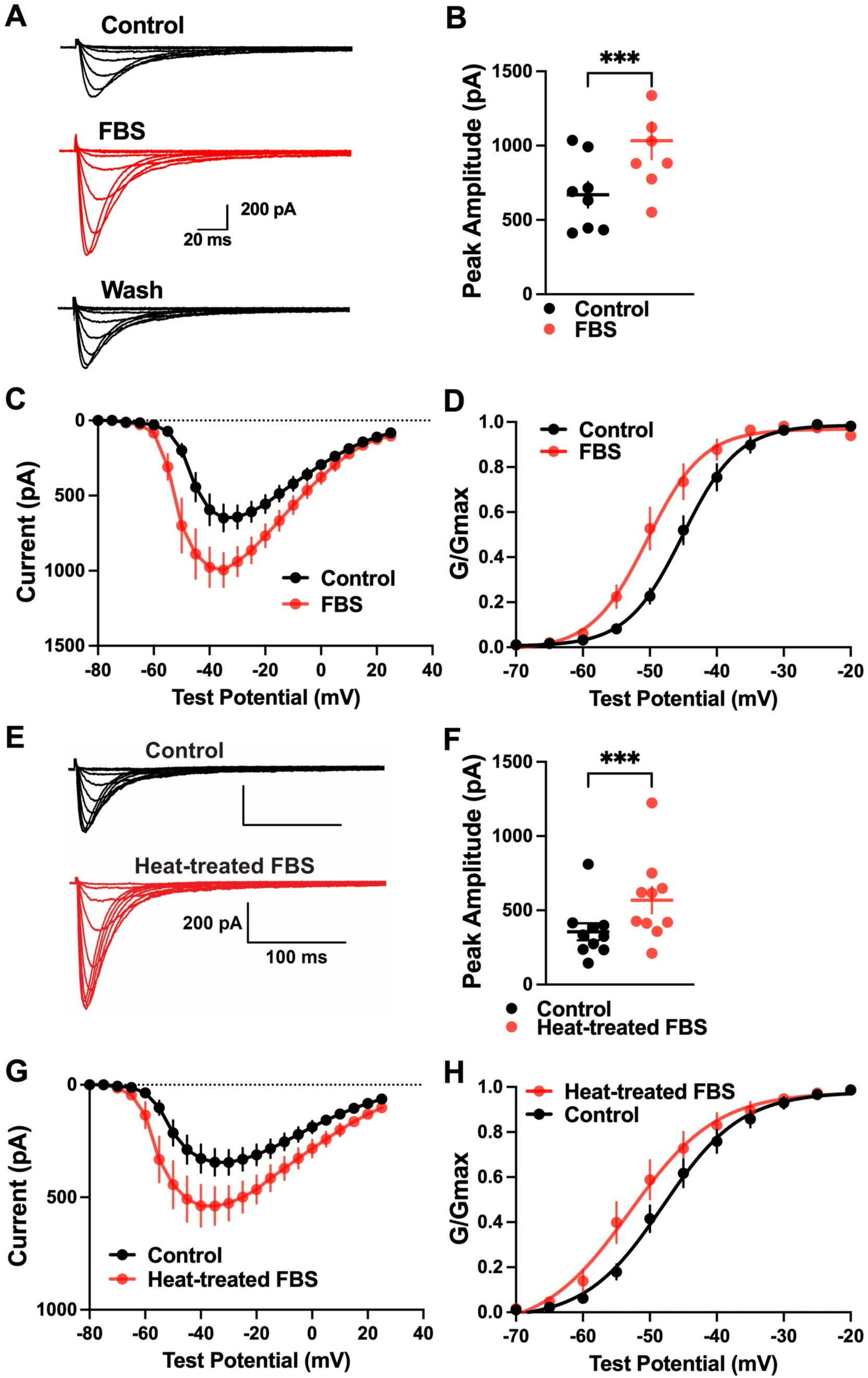
Fetal bovine serum (FBS) reversibly increases current amplitude and activation kinetics of recombinant human Ca_V_3.2 currents in a heat-resistant manner. **(A)** Families of inward calcium current traces recorded from Ca_V_3.2 channel-expressing HEK- 293 cells before (black) and after perfusion with 1% FBS (red), as well as after washing with external solution (black) (V*h* = -90 mV, V*t* = -70 mV through -30 mV). **(B)** Maximal peak current amplitudes shown as dot-plots (mean peak current amplitude at baseline: 669.3 ± 85.9 pA, mean amplitude in the presence of 1% FBS: 1033.0 ± 124.6 pA, n=8, p<0.001, two-tailed paired t-test). **(C)** Averaged current-voltage (IV) curve shows that FBS (red symbols) increased the amplitudes of peak currents compared with control pre-drug conditions (black symbols) over a wide range of test potentials (n=8). **(D)** Treatment of HEK-293 cells with 1% FBS resulted in a ∼5 mV hyperpolarizing shift in V_50_ of Ca_V_3.2 channel activation as determined using the Boltzmann function (V_50_ control = -44.69 ± 1.11 mV, V_50_ serum = -49.62 ± 1.54 mV, n=8; p<0.001, two-tailed paired t-test). **(E)** Examples of original current traces before (black) and after perfusion with 1% heat-treated FBS (red) at different test potentials (V*h* = -90 mV, V*t* = -70 mV through -30 mV). **(F)** Maximal peak current amplitudes shown as dotted plots (mean peak current amplitude for baseline: 355.8 ± 57.2 pA, mean amplitude for heat-treated FBS: 568.6 ± 88.9 pA, n=10, p<0.001, two-tailed paired t-test). **(G)** Averaged IV curve shows that heat-treated serum (red symbols) maintains its ability to increase the amplitudes of peak currents compared with control baseline conditions (black symbols) over a wide range of test potentials (n=10). **(H)** Treatment of HEK-293 cells with 1% heat-treated FBS resulted in an approximately 4-mV hyperpolarizing shift of the activation curve (V_50_ control = -47.40 ± 1.45 mV, V_50_ serum = -51.59 ± 2.16 mV, n=10; p<0.005, two-tailed paired t-test). Bars on panels A and E indicate calibration. Significance on all panels: * p<0.05; ** p<0.01; *** p<0.001.

We next subjected FBS to heat treatment by boiling it for 10–15 min. Figure 1 E-H shows that the T-current augmenting properties of FBS were well preserved after heat treatment, which is expected to inactivate all large proteins.

### Serum enhances the excitability of rat DRG nociceptors *in vitro*

To probe the effects of serum on T-currents and I_Na+_ in the same experiment, we recorded IV curves in external Tyrode’s solution by applying depolarizing steps in 2-mV increments. Composite inward currents in Figure 2A depict this experiment in a representative smaller rat DRG neuron. We first recorded subthreshold inward currents under baseline conditions (Figure 2A, top panel). Then, we added 1% FBS to the external solution for 5 min and observed a larger and faster inactivating inward component, representing I_Na+_, along with the slower inactivating component (black arrow), representing the T-current (Figure 2A, middle panel). We next co-applied a T-channel blocker, 3β-OH [(3*β*,5*β*,17*β*)-3-hydroxyandrostane-17-carbonitrile] ^19,20^ together with FBS. Traces in Figure 2A (bottom) show that only the subthreshold inward currents were recorded in the presence of 1% FBS and 3*β*-OH, the same as in the baseline conditions. The summary bar graph in Figure 2B from 9 DRG neurons shows that 1% FBS significantly decreased the average activation threshold by -8.4 ± 1.1 mV (black solid bar) compared with the baseline (solid horizontal line). In contrast, when FBS and 3*β*-OH were co-applied on the same neurons (open bar), the average activation threshold was not significantly different from the control baseline values. This finding was independently confirmed with another T-channel inhibitor such as (3,5-dichloro-N-[1-(2,2-dimethyl-tetrahydro-pyran-4-ylmethyl)-4-fluoro-piperidin-4- ylmethyl]-benzamide (TTA-P2) ^21^ at 3 μM (n=7, data not shown). Furthermore, we previously reported that presynaptic Ca_V_3.2 channels regulate glutamate release in dorsal horn (DH) neurons of the spinal cord, a major pain processing region ^9^. Hence, we hypothesized that FBS also promotes excitability in the DH neurons. For these experiments, we recorded spontaneous excitatory postsynaptic currents (sEPSCs) from the superficial lamina I DH neurons of adult rats and found that 1% FBS increased the frequency of sEPSCs by two-fold, and this effect was fully reversed by 1 μM TTA-P2 (Supplementary Figure 3). Overall, these data demonstrate a crucial role for T-channels in the mechanisms underlying serum-induced hyperexcitability of both peripheral and central pain pathways.

**Figure 2.**
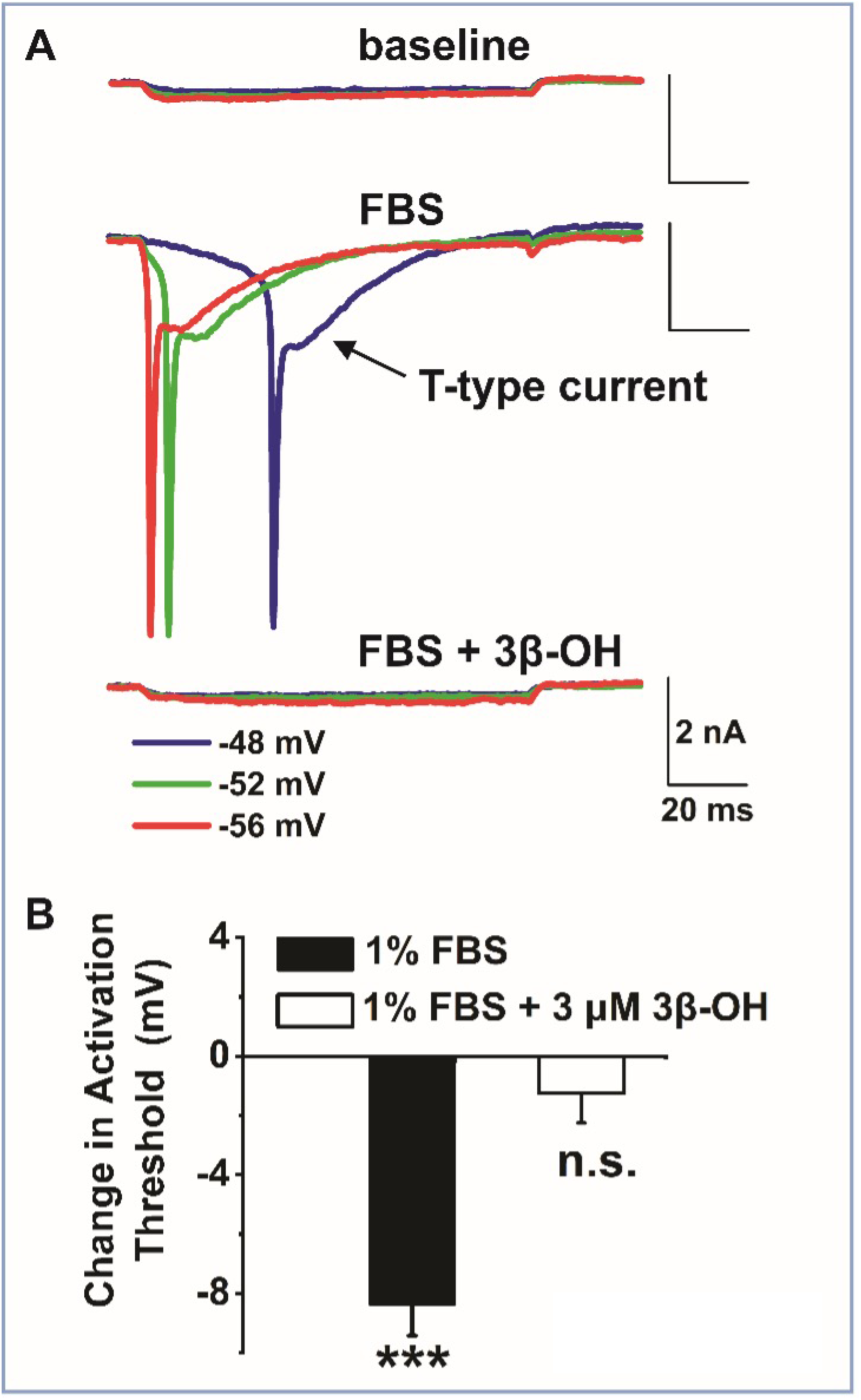
Serum facilitates T-channel-dependent excitability of putative nociceptive DRG neurons from adult rats. **(A)** Original traces from a representative DRG neuron show that minimal inward currents could be elicited at -48, -52, and -56 mV in baseline conditions. After perfusion with 1% FBS, the voltage steps at -48, -52, and -56 mV resulted in faster inactivating voltage-gated sodium currents and more slowly inactivating T-type currents (black arrow). Perfusion with 1% FBS in the presence of a selective T-channel blocker 3β-OH (3 μM) abolished the current activation at - 48, -52, and -56 mV, and only minimal inward currents were recorded, similar to baseline pre- drug conditions. (V*h* -90 mV, V*t* -60 to -20 mV in 4-mV increments). **(B)** Bar graph shows the average data from 9 DRG neurons following the same experiment as in panel A. Note that FBS decreased the average threshold for sodium current activation by - 8.4 ± 1.1 mV (black-filled bar) compared with the baseline in the same DRG neurons (p<0.001, two-tailed paired t-test). In contrast, when FBS and 3β-OH were co-applied on the same neurons (open bar), the average threshold for activation was not significantly different from the baseline (-1.2 ± 0.8 mV, p=0.3, two-tailed paired t-test).

### Serum induces peripheral heat hyperalgesia *in vivo*

We next examined if serum facilitates pain transmission in live animals. Thermal nociception to noxious heat stimuli was evaluated in rats before and after intraplantar (i.pl.) injection with 100 μL of 1% or 3% FBS into the right hind paw by measuring paw withdrawal latencies (PWLs) to noxious heat stimuli (Figure 3A). To exclude the possibility that the tissue trauma incurred during injection was responsible for changes in nociception, control rats were given i.pl. injections of vehicle and evaluated using the same protocol. As expected, control animals exhibited no right-left difference in PWLs after injection of vehicle (Figure 3B). In the experimental group, injections of 1% FBS (Figure 3C) and 3% FBS (Figure 3D) resulted in a significant, although transient, decrease in right paw thermal PWLs by approximately 20% and 40%, respectively, at 10, 20, and 60 min post-injection in comparison to uninjected control paws, indicating the development of prominent serum-induced heat hyperalgesia. Next, we co-injected 30 nM 3β-OH along with 3% serum. Thermal PWLs did not deviate from the baseline values upon co-injection, indicating that serum-induced thermal hyperalgesia in rats likely involves peripheral Ca_V_3.2 channels (Figure 3E).

**Figure 3.**
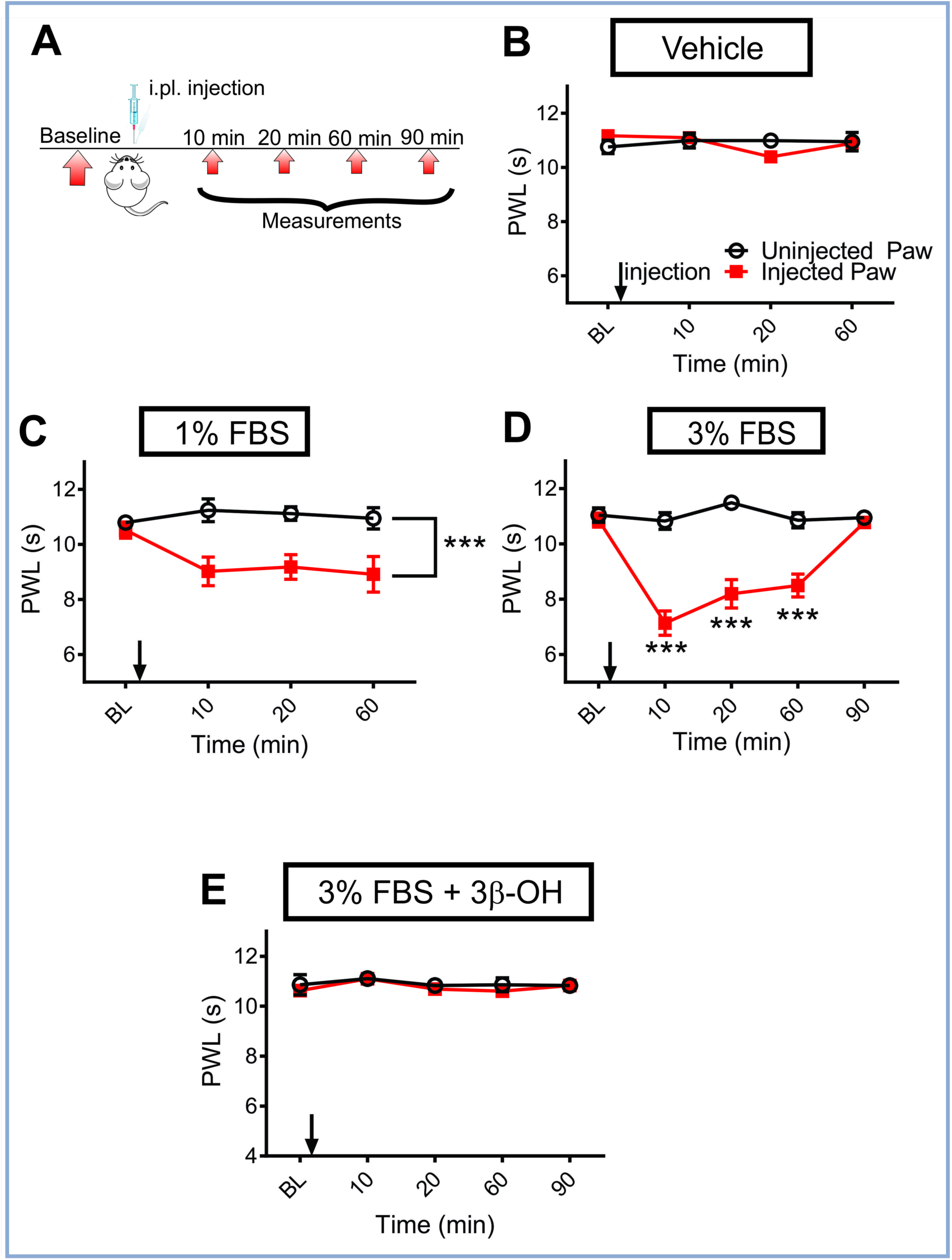
Thermal heat hyperalgesia in rats induced by intraplantar (i.pl.) injections of FBS is T-channel dependent. **(A)** Time course of the experiment showing pre-injection baseline measurements of responses to a thermal heat stimulus determined 2 days before the experiment. Heat nociception was assessed 10, 20, 60, and 90 min after i.pl. injection of either vehicle (as denoted by arrows), FBS, and/or the selective T-channel blocker 3β-OH. **(B)** Lack of effect of i.pl. vehicle injections on heat nociception in rats compared with the uninjected paw (n=6 rats, two-way repeated measures ANOVA; p=0.84). **(C)** Analysis of the hyperalgesic effect of i.pl. injections of 1% FBS on heat nociception in rats compared with the uninjected paw (n=6 rats; two-way repeated measures ANOVA; treatment: F(1,10)=24.13, p=0.0006). **(D)** Analysis of the effect of i.pl. injections of 3% FBS on heat nociception in rats compared with the uninjected paw (n=6 rats; two-way repeated measures ANOVA with Bonferroni’s multiple comparison test; interaction: F(4,25)=24.95, p<0.0001; treatment: F(1,25)=166.7, p<0.0001). ***p<0.001. **(E)** Analysis of the antinociceptive effect of the i.pl. injection of 30 nM 3β-OH on heat hyperalgesia evoked with 3% FBS in rats, as compared with the noninjected paw (n=6 rats; two-way repeated measures ANOVA; treatment: p=0.41). Each data point represents the average thresholds for paw withdrawal latency (PWL) ± SEM.

### Ca_V_3.2 channels in peripheral sensory neurons are required for serum-induced hypersensitivity to mechanical stimuli *in vivo*

We next utilized mouse genetics and assayed mechanical sensitivity to further validate the notion that Ca_V_3.2 channels are important for the pro-nociceptive effects of serum. We measured baseline thresholds for mechanical paw withdrawal responses (PWRs) in both hind paws before i.pl. injection (baseline) and at 10, 20, 30, 60, and 90 min following injections of 1% FBS or vehicle (normal saline), as depicted in Figure 4A. Vehicle injections had no effect on PWRs in WT mice (gray symbols, Figure 4B). On the other hand, serum-induced mechanical hypersensitivity in WT mice was evident by a reduction in the PWR threshold of approximately 50% after 10 min, 40% after 20 min, and 20% after 30 min post injections (red symbols, Figure 4B). Thresholds for PWRs in the uninjected (left) paws remained stable throughout the testing period, indicating the lack of a systemic effect of FBS (open symbols, Figure 4B). In contrast to FBS, we found that i.pl. injections of vehicle (VEH, gray symbols in Figure 4B) did not significantly affect baseline PWRs in WT mice. Importantly, i.pl. injections of FBS had no effect on the PWR thresholds in Ca_V_3.2 knock-out (KO) mice (red symbols, Figure 4C) compared with the uninjected paws (open symbols, Figure 4C). As reported previously ^22–24^, baseline mechanical sensitivities are similar between Ca_V_3.2 null mice and their WT counterparts. Because only WT mice exhibited decreased thresholds for PWRs after i.pl. injections of FBS, however, we conclude that Ca_V_3.2 channels in peripheral sensory neurons are required for serum-induced hypersensitivity to mechanical stimuli *in vivo*.

**Figure 4.**
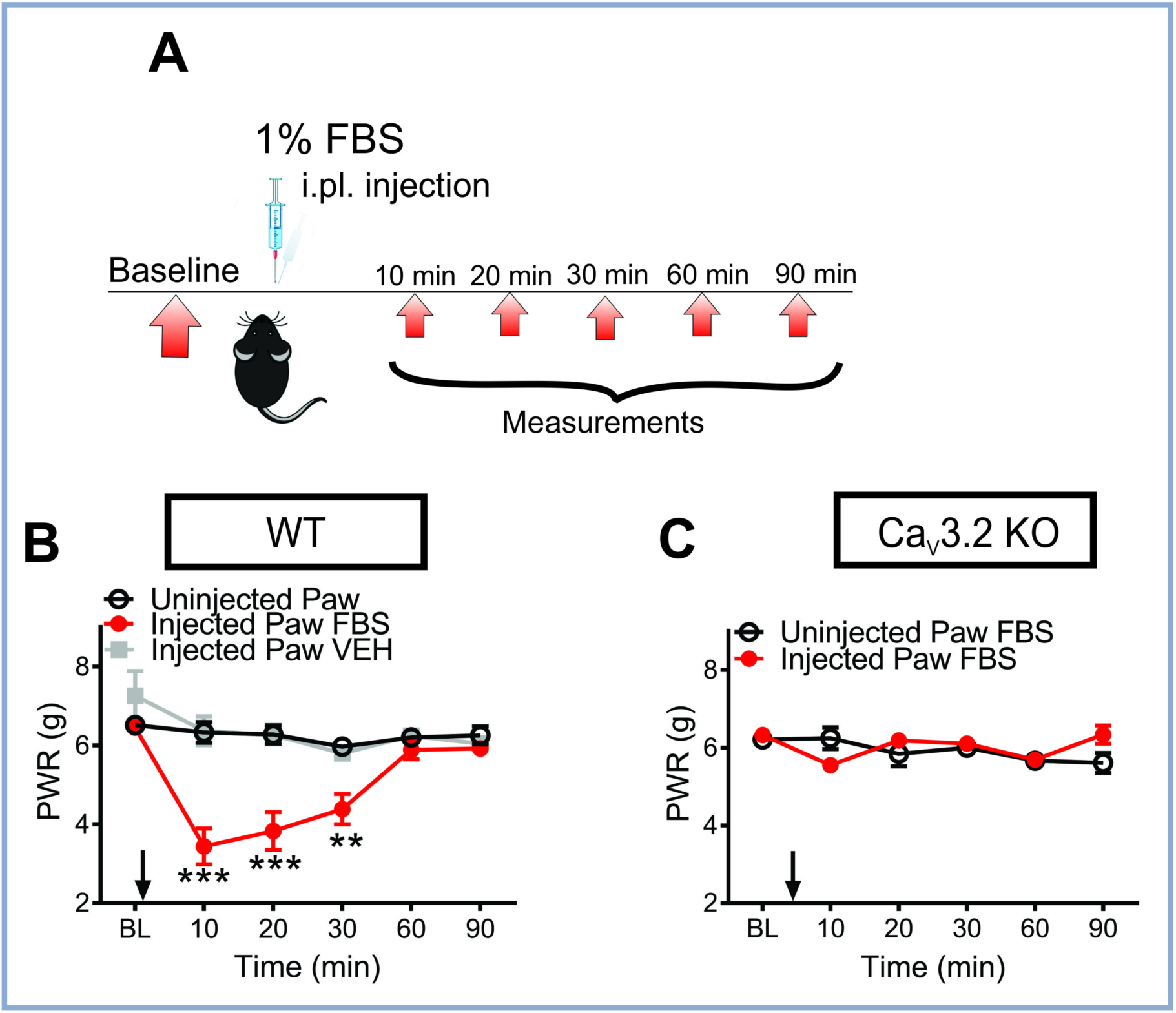
The Ca_V_3.2 channel in peripheral sensory neurons is required for serum-induced hypersensitivity to mechanical stimuli *in vivo*. **(A)** Time course showing the pre-injection baseline measurements of responses to mechanical stimulus in the WT female mice determined 2 days before the experiment. Mechanical sensitivity was assessed 10, 20, 30, 60, and 90 min after i.pl. injection of the dialyzed 1% FBS as denoted by arrows. **(B)** Analysis of the hyperalgesic effect of i.pl. injection of FBS on mechanical sensitivity to Von Frey filament in the WT mice as compared with the vehicle group (n=8-9 per group; two-way repeated-measures ANOVA with Sidak’s multiple comparison test; interaction: F(10,155) =6.78, p<0.0001; treatment: F(2,31)=20.18, p<0.0001). **(C)** The graph shows the lack of an effect of i.pl. injection of FBS on mechanical sensitivity to Von Frey filament in Ca_V_3.2 KO female mice as compared with the uninjected paw (n=12 per group; two-way repeated measures ANOVA; treatment: p=0.59). Each data point represents the average thresholds for the paw withdrawal response (PWR) ± SEM.

### Serum enhances Ca_V_3.2 currents *via* trace metal chelation of an H191 high-affinity histidine residue

Next, we examined the effects of FBS on peak T-currents (V*h* -90 mV, V*t* -40 mV) compared with HVA calcium currents (V*h* -55 mV, V*t* 0 mV) in smaller rat DRG neurons. Original traces from a representative DRG neuron presented in Figures 5A and 5B depict the effects of 1% FBS on T-type and HVA currents, respectively. While the T-current is reversibly enhanced (∼two-fold) as expected, the HVA current was not significantly affected in these same DRG neurons (n=3, 90±10% of baseline current, p=0.2). This selective increase in T-current vs. HVA current was mirrored in our experiments using human serum and human DRG cells (Supplemental Figure 2). Similarly, our previous studies established that only DRG T-currents, and not HVA currents, are augmented by reducing agents such as dithiothreitol (DTT) and L-cysteine (L-cys) and diminished with the oxidizing agent 5,5’-dithio-bis-(2-nitrobenzoic acid, DTNB) ^7,25,26^. To determine whether serum potentiates T-currents v*ia* a redox-specific mechanism, serum was co-applied with DTNB. The representative time course from the experiment depicted in Figure 5C shows that peak amplitudes of DRG T-currents observed after co-application of 1% FBS and DTT were not larger than those elicited after adding either agent alone. Accordingly, we hypothesized that the active component of serum may include amino acids such as cysteine and histidine that are involved in redox and/or metal chelating reactions ^27–30^. Hence, we recorded T-currents when 1% FBS was co-applied with a strong oxidizing agent DTNB. DTNB inactivates free cysteine groups in proteins by oxidizing them and forming covalent disulfide bonds, but is unlikely to affect histidine groups on the proteins. Co-application of 1% FBS with 100 μM DTNB still increased T-current amplitudes (∼two-fold), which was not different from the change in current with 1% FBS alone in these neurons (Figure 5D). As expected, when L-Cys was applied to the same DRG neurons, it increased the current amplitude like FBS, but its effect on T-currents was completely abolished in the presence of DTNB (Figure 5D). Overall, DTNB failed to inhibit T-current increase by 1% FBS (n=4), while it completely abolished response to L-Cys in the same cells (n=3). These data strongly suggest that putative histidine groups rather than cysteine groups are involved in mediating the effects of FBS on Ca_V_3.2 T-currents.

**Figure 5:**
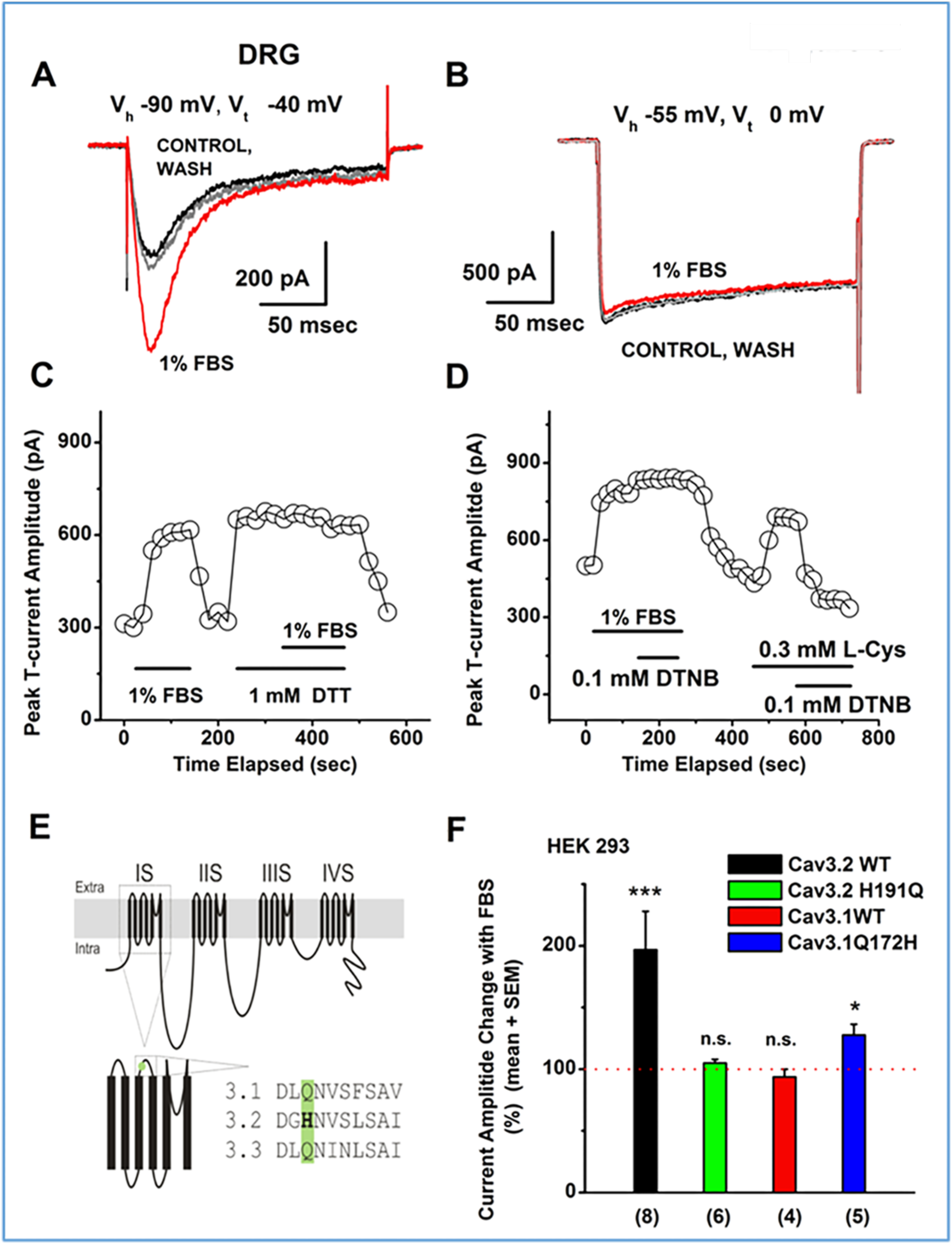
Serum potentiates T-currents *via* trace metal chelation of a critical Ca_V_3.2 channel histidine residue. **(A)** Traces of T-currents (V*h* -90 mV, V*t* -30 mV) in a representative rat DRG neuron before (black trace) and after wash (gray traces), as well as during bath application of 1% serum (FBS, red trace), which reversibly increased the peak inward current by approximately two-fold. **(B)** Traces of high-voltage-activated (HVA) calcium currents from the same rat DRG cell (V*h* -55 mV, V*t* 0 mV) depicted on panel A before (black trace), during the bath application of 1% FBS (red trace), and after wash (gray trace). Note that FBS had little effect on the inward current under these recording conditions. **(C)** Time course from a representative experiment in a rat DRG neuron shows peak T-current amplitudes (V*h* -90 mV, V*t* -30 mV) during applications of 1 mM DTT and 1% FBS. Initial application of FBS alone reversibly increased the T-current amplitude by approximately two-fold. After returning the T-current amplitude to the baseline level by washing out the FBS with external bath solution, DTT was applied alone before FBS and DTT were co-applied. Note that the application of FBS had no additional effect on the current amplitude after pretreatment with DTT, suggesting a similar mechanism of action. **(D)** Time course from another experiment in a rat DRG neuron shows peak T-current amplitudes (V*h* -90 mV, V*t* -30 mV) during applications of 1% FBS, 0.1 mM DTNB, and 0.3 mM L-cysteine (L-Cys). Initial application of FBS alone reversibly increased the T-current amplitude from 450 pA to 850 pA and this effect was not reversed with the co-application of DTNB. After return of the T- current amplitude to the baseline level, L-Cys was applied alone and then L-Cys and DTNB were sequentially co-applied. Note that the co-application of DTNB completely reversed the increase in the T-current amplitude induced by the application of L-Cys alone. **(E)** A diagram of the Ca_V_3.2 channel depicts that the metal-binding histidine residue at position 191 is unique to the Ca_V_3.2 isoform of T-type calcium channels. **(F)** Bar graph from multiple experiments shows that FBS potentiates amplitudes of recombinant Ca_V_3.2 currents in HEK-293 cells by 97 ± 31% but exerts no significant effect on recombinant WT Ca_V_3.1 or mutated H191Q Ca_V_3.2 currents, both of which lack the critical H191 histidine residue. When the critical histidine was introduced to Ca_V_3.1 channels (Ca_V_3.1 Q172H), the application of FBS significantly increased the T-current amplitude in these mutated channels by 28 ± 9%. Red dashed line indicates normalized baseline T-current amplitudes. Number of cells in each recording condition is indicated in parenthesis. Asterisks denote significance by two-tailed paired Student t-test : *, p<0.05; ***, p<0.001; n.s., p>0.05.

Previous molecular studies demonstrated that substitution of a unique extracellular histidine residue with glutamine (H191Q) completely abolished Ca_V_3.2 current inhibition by trace metals such as zinc, copper, and nickel ^26,31^. Figure 5E depicts a scheme of H191 in the Ca_V_3.2 isoform compared with Ca_V_3.1 and Ca_V_3.3 isoforms. Next, we recorded currents from WT Ca_V_3.2 channels, WT Ca_V_3.1 channels, and Ca_V_3.2 H191Q-mutant channels expressed in HEK-293 cells before and after treatment with 1% FBS (Figure 5F). As expected, 1% FBS robustly enhanced currents in WT Ca_V_3.2 channels (black bar) by approximately two-fold but not in Ca_V_3.2-H191Q (green bar) and WT Ca_V_3.1 channels (red bar). This strongly suggests that serum worked at least in part by relieving the tonic inhibition of Ca_V_3.2 channels by chelating trace metals from the H191 residue. We next attempted to reverse the serum insensitivity of Ca_V_3.1 channels by performing the analogous reverse mutation (Q172H). We observed an approximately 30% enhancement of Ca_V_3.1Q171H currents in the presence of 1% FBS compared with baseline (blue bar, Figure 5F). Taken together, these findings indicate that serum contains at least one molecule capable of influencing Ca_V_3.2 channel properties by trace metal chelation at the critical histidine residue rather than by redox-dependent mechanisms such as the modification of cysteine residues.

### Effects of serum on mechanical hypersensitivity *in vivo* are abolished in Ca_V_3.2 H191Q KI mice

Next, we generated a Ca_V_3.2 H191Q KI mouse (Supplementary Figure 4) and performed behavioral pain experiments to evaluate the effects of this point mutation on both peripheral and central effects of FBS on mechanical sensitivity. As expected, i.pl. injection of 1% FBS induced reversible decreases in PWRs in ipsilateral paws of WT mice at 10, 20, and 40 min following the injections (red symbols, Figure 6A) compared with the contralateral non-injected paws (dark blue symbols, Figure 6A). Importantly, Figure 6B shows that i.pl. injections of 1% FBS had no effect on mechanical PWRs in the ipsilateral paws of Ca_V_3.2 H191Q KI mice (red symbols) compared with the contralateral paws (dark blue symbols). In contrast to FBS, we found that i.pl. injections of vehicle did not significantly affect baseline PWRs in WT mice (data not shown).

**Figure 6.**
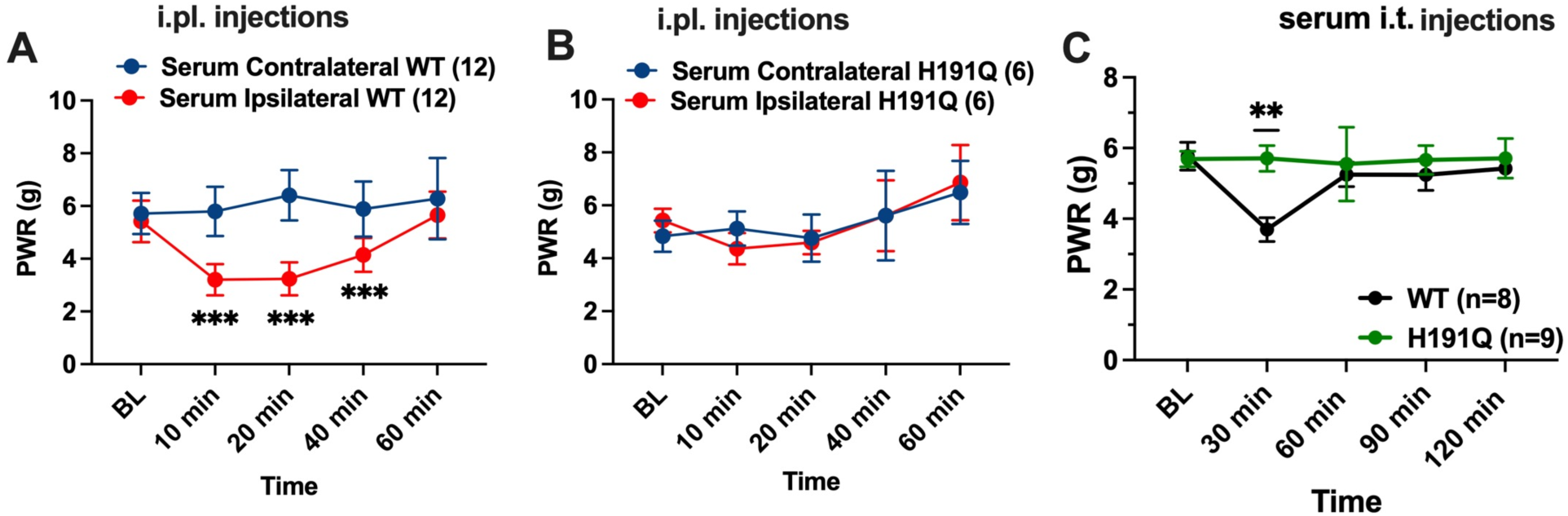
The H191 residue of Ca_V_3.2 channel in peripheral and central sensory neurons is required for serum-induced hypersensitivity to mechanical stimuli *in vivo*. Analysis of the effect of the i.pl. FBS injection in female WT (**A**) and H191Q knock-in mice (**B**) on mechanical hypersensitivity; n=6-12 mice per group. Two-way repeated measures ANOVA: Interaction F_(4,44)_=11.08, p<0.001; Time F_(4,44)_=11, p<0.001; Injection F_(1,11)_=175.3, p<0.001, post-hoc presented on Figure. (**C**) Time course of the effect of serum i.t. injected in WT and knock-in mice on mechanical hypersensitivity, n=8,9 mice. Two-way RM ANOVA: Interaction F_(4,60)_=1.53, p=0.204; Time F_(2.2,33)_=1.42, p=0.256; Genotype F_(1,15)_=2.21, p=0.157; post hoc presented on Figure. Each data point represents the mean ± SEM.” Asterisks denote significance: **, p<0.01; ***, p<0.001;

We next used intrathecal (i.t.) injections of FBS to compare its effects on spinal nociception in WT and Ca_V_3.2 H191Q KI littermates. For these experiments, we first determined baseline mechanical PWRs on both hind paws. We then injected 10 μL of 3% FBS i.t. at the L4-L6 spinal region under isoflurane anesthesia. Mice were allowed to recover fully from anesthesia and PWRs in both hind paws were measured again. Average PWR values at 30-, 60-, and 90-min post-injection are summarized in Figure 6C. At the 30-min time point, the average PWRs were decreased by approximately 40% from baseline in WT mice, indicating prominent hypersensitivity to mechanical stimuli, with a return to baseline values at the 60- and 90-min time points. In contrast, PWRs remained unchanged throughout the 120-min testing period in the KI group. Injections of vehicle (saline) did not affect PWRs in either of these cohorts (data not shown). These data strongly suggest that the H191 residue of the Ca_V_3.2 channel is required for serum-induced hypersensitivity to mechanical stimuli in both peripheral and central pain pathways.

### Partial purification and size-exclusion dialysis indicate that the T-channel modulating substance is likely a heat-stable, acid-stable, hydrophilic peptide with a low MW

Next, we performed an initial purification of FBS using acutely dissociated rat DRG neurons to characterize the properties of the T-channel modulating compound, as summarized in Table 1 (Supplementary Figure 5). Consistent with our experiments using recombinant Ca_V_3.2 channels (Figure 1), we found that a 1% heat-treated fraction of FBS still retained robust potentiating activity on native T-currents (203 ± 26%, n=3, p<0.0001). We also found that incubating serum overnight at a pH <2 did not significantly affect its effect (176 ± 7%, n=5, p<0.0001). In contrast, when FBS was incubated overnight at pH >10, the effect was completely abolished (107 ± 5%, n=5, p=0.9995). In addition, 10 μM copper in external solution also completely abolished the effect of 1% FBS on DRG T-currents (112 ± 21%, n=7, p=0.9576). To test whether the active substance is hydrophilic, we separated hydrophobic and hydrophilic components by treating FBS with 10% chloroform. The hydrophilic component retained its full potentiating effect on T-currents as shown on Table 1 (240 ± 24%, n=4, p<0.0001), while the hydrophobic component was ineffective (data not shown).

We subsequently determined the approximate MW of this T-channel modulator by screening its effects on recombinant Ca_V_3.2 channels and using FBS dialyzed through filters with different MW cutoffs. Ca_V_3.2 currents were reliably augmented by serum fractions in the range of 3–5 kDa (n=6, Supplementary Figure 6) and 3–7 kDa (n=5 data not shown), but not by serum fractions in the range of 5–7 kDa (n=5, data not shown). Next, we proceeded with tandem mass spectrometry analysis of our serum samples.

### Proteomic analysis of the FBS reveals an active peptide: albumin (1-26)

Proteomic analysis of the active fraction of partially purified FBS using dialysis filters with different MW cutoffs revealed approximately 30 peptides, with the 10 most abundant peptides listed in Table 2 (Supplementary Figure 7). Using their amino acid sequences as a template, we synthesized a histidine-rich bovine albumin N-terminal peptide with MW of ∼3 kDa (albumin 1-26, DTHKSEIAHRFKDLGEEHFKGLVLIA) to determine whether this peptide with three histidine residues (shaded in yellow) possesses the metal-chelating property needed to disinhibit Ca_V_3.2 channels. Indeed, albumin (1-26) fragment displayed electrophysiological properties closely mimicking those observed with FBS (Figure 7). Representative traces from HEK-293 cells before (black line) and after the application of albumin (1-26) (red line), which increased Ca_V_3.2 current amplitudes approximately two-fold are depicted in Figure 7A. Average peak current amplitudes showed a robust increase when perfused with albumin (1-26) peptide (Figure 7B). In addition, 100 μM albumin (1-26) peptide induced an up to two-fold increase in T-current amplitudes across a wide range of test potentials (-55 mV to -5 mV) (Figure 7C) and induced a significant hyperpolarizing shift (∼3 mV) in channel gating (Figure 7D). Importantly, Figure 7E-H shows that the facilitatory effect of albumin (1-26) peptide on Ca_V_3.2 currents is preserved after heat treatment. As expected, physiological concentrations of full sequence bovine serum albumin (BSA) protein also robustly increased Ca_V_3.2 current densities and facilitated channel gating, but heat treatment completely abolished the effects (Supplementary Figure 8). We next investigated whether T-current augmentation by bovine albumin (1-26) peptide is conserved in other mammalian species. Peptides with corresponding sequences from human, horse, mouse, and rat all increased Ca_V_3.2 currents in stably transfected HEK-293 cells (Supplementary Figure 9). Table 3 summarizes the amino acid sequences of these peptides (Supplementary Figure 10).

**Figure 7.**
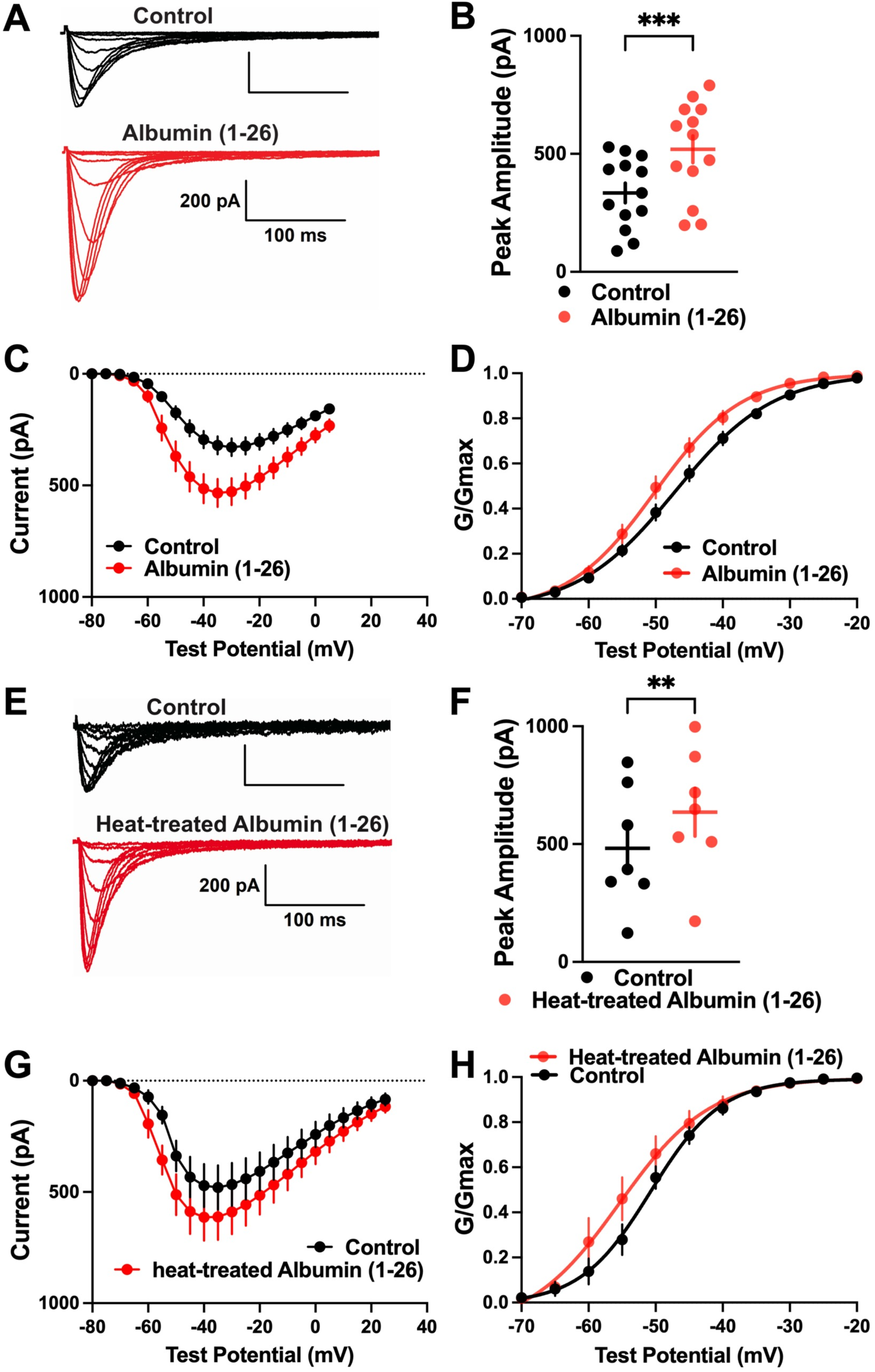
The albumin (1-26) acts as a novel endogenous Ca_V_3.2 channel modulator. **(A)** Representative Ca_V_3.2 T-current traces before (black) and after the addition of 0.1 mM albumin (1-26) (red) to the external solution in HEK-293 cells expressing recombinant Ca_V_3.2 channels (V*h* = -90 mV, V*t* = -70 mV through -30 mV). **(B)** Comparison of the maximal peak current amplitudes displayed as dot-plots (mean peak current amplitude at baseline: 334.6 ± 42.0 pA, mean peak current amplitude after addition of 0.1 mM albumin (1-26) peptide: 519.6 ± 56.4 pA, n=13, p<0.001, two-tailed paired t-test). **(C)** Graph displaying the current-voltage relationship of activation showing the increase in current amplitudes across a wide voltage spectrum in the presence of albumin (1-26) peptide. **(D)** Voltage-dependent activation kinetics show a significant leftward shift after fitting with the Boltzmann function under the influence of 0.1 mM albumin (1-26) peptide (baseline V_50_: -47.24 ± 1.02 mV compared with albumin (1-26) V_50_: -50.08 ± 1.21 mV, n=17, p=0.003, two-tailed paired Student’s t-test). **(E)** Representative Ca_V_3.2 current traces of current-voltage relationship before (black) and after (red) the addition of 0.1 mM heat-treated albumin (1-26) peptide to the external solution (V*h* = - 90 mV, V*t* = -70 mV through -30 mV). **(F)** Dot-plot depiction of the maximal peak current amplitudes before and after addition of the 0.1 mM heat-treated albumin (1-26) fragment shows the maintained ability to significantly augment the Ca_V_3.2 currents (control: 482.4 ± 97.7 pA, heat-treated peptide: 635.4 ± 101.8 pA, n=7, p=0.003, two-tailed paired t-test). **(G)** Graph displaying the current-voltage relationship of activation showing the maintained increase in current amplitudes across the voltage spectrum after heat treatment of the albumin (1-26) peptide. **(H)** Voltage-dependent activation curves show a leftward shift of the half activation voltage V_50_ from -49.1 ± 1.2 mV in baseline conditions to -53.4 ± 1.5 mV after the application of heat-treated albumin (1-26) (n=7, p=0.02, two-tailed paired t-test).

### Albumin 1-26 induced mechanical hyperalgesia in rats in a T-channel-dependent manner

In the following *in vivo* study, i.pl. injections of albumin (1-26), but not injections of vehicle, into rat hind paws led to an approximately 30% decrease in the PWR at 10 and 20 min post-injection (Figure 8A-B). Importantly, co-injectons of 3β-OH completely abolished this effect (Figure 8C) but 3β-OH alone had no clear effect on the PWR threshold (Figure 8D). Injections of BSA, however, resulted in an approximately 50% decrease in PWR thresholds at 10, 20, and 40 min after injection (Figure 8E). As with albumin (1-26), BSA lost its hyperalgesia-inducing property when co-applied with 3β-OH suggesting a role of T-channels (Figure 8F).

**Figure 8.**
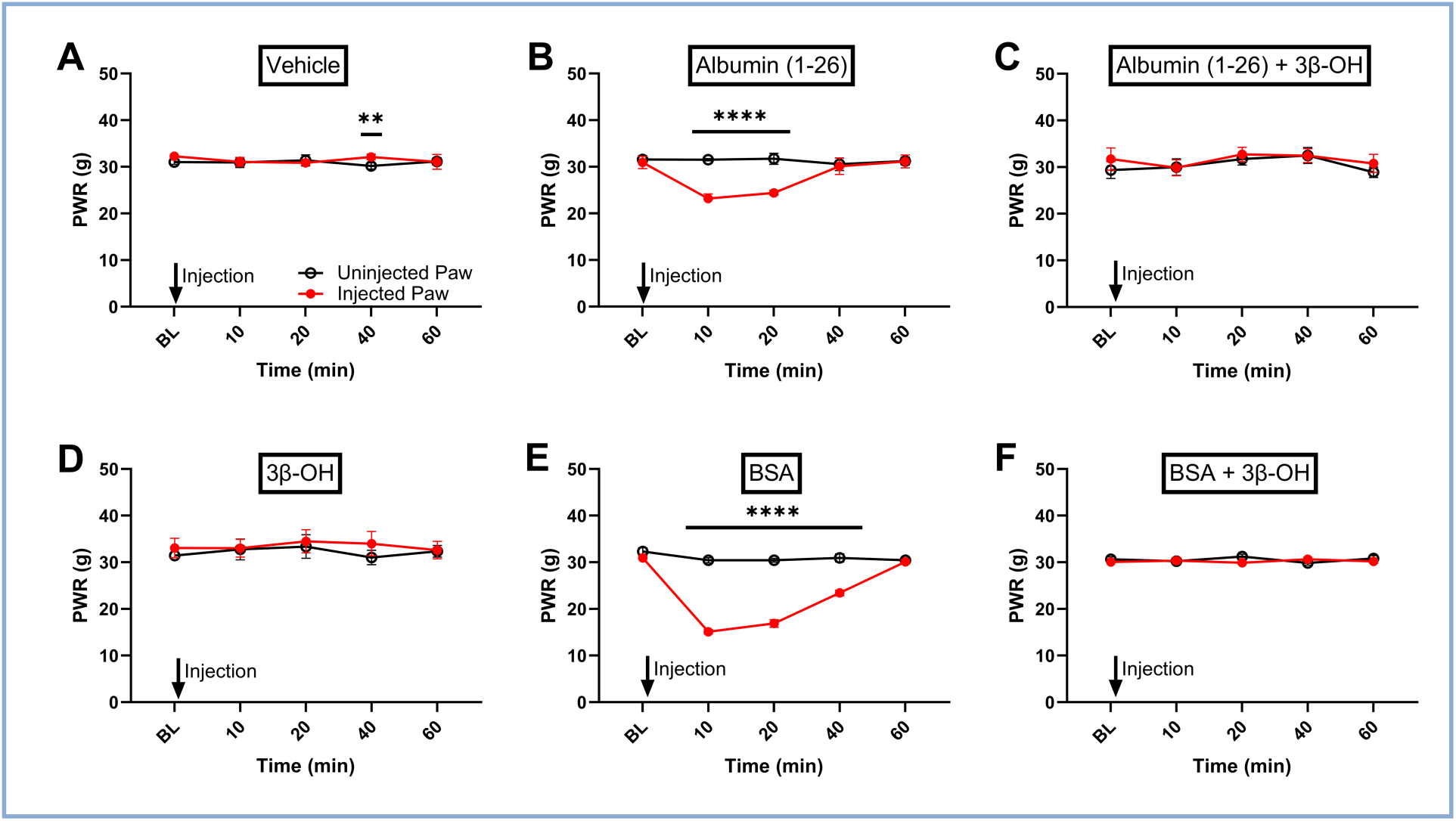
Paw injections of bovine serum albumin (BSA) and N-terminal bovine albumin (1-26) peptide induce T-channel-dependent mechanical hypersensitivity in rats. **(A)** Paw withdrawal response (PWR) thresholds in paws injected with saline (red) compared with the contralateral paws at different time points after injections of vehicle were almost overlapping, but showed a very small difference at 40 min after injection (n=6 in each group, two-way repeated-measure ANOVA with Sidak’s multiple comparison’s test, treatment: F(1,5)=11.39, p=0.0198, interaction: F(4,20)=3.71, p=0.02, Sidak’s: at 40 min after injection: p=0.0075). **(B)** Graph showing a significant reversible decrease of the PWR threshold in rats after i.pl. injection of 0.2 mM bovine albumin (1-26) peptide of about 33% (n=6 per group, two-way repeated-measure ANOVA with Sidak’s multiple comparisons test, treatment: F(1,5)=177.5, p<0.0001; interaction: F(4,20)=33.81, p<0.0001; Sidak’s: p<0.0001 at 10 and 20 min). This effect was completely abolished when bovine albumin (1-26) peptide was co-applied with the T-type channel blocker 3*β*-OH (n=6, treatment: F(1,5)=0.58, p=0.48; interaction: F(4,20)=0.27, p=0.89) **(C)**. **(D)** Graph showing the lack of difference between the PWRs in the injected vs. the uninjected paw, when applying 3*β*-OH alone (n=6, treatment: F(1,5)=1.07, p=0.35; interaction: F(4,20)=0.18, p=0.95). **(E)** Intraplantar injections of 0.2 mM bovine serum albumin (BSA) alone induced robust reversible hyperalgesia (n=6 in each group, two-way repeated-measure ANOVA followed by Sidak’s multiple comparison, treatment: F(1,5)=523.5, p<0.0001; interaction: F(4,20)=78.2, p<0.0001, Sidak’s: p<0.0001 at 10, 20, and 40 min). **(F)** This effect could also be completely abolished with the addition of the T-type channel blocker 3*β*-OH (n=6, treatment: F(1,5)=1.09, p=0.34; interaction: F(4,20)=1.7, p=0.19).

## Discussion

Here, we described for the first time the effects of mammalian sera from different species and an endogenous albumin-related peptide on the biophysical properties of Ca_V_3.2 channels *in vitro* and on nociceptive-like behavior in animal models *in vivo*. We showed that serum is a selective and potent potentiator of both native and recombinant human Ca_V_3.2 currents by removing tonic inhibition induced by trace metals. Further, our findings demonstrate that even 100-fold diluted serum solutions can profoundly enhance the amplitude of native and recombinant Ca_V_3.2 T-current *in vitro* and directly sensitize peripheral and central nociceptors to thermal and mechanical stimuli in WT rats and mice. In contrast, injections of serum in Ca_V_3.2 KO mice and Ca_V_3.2 H191Q KI mice did not affect baseline responses to mechanical stimuli. These data indicate that Ca_V_3.2 channels and their specific metal-binding H191 residue are required for sensitization of both peripheral and spinal nociceptors by serum. Taken together, these findings indicate that serum contains at least one molecule capable of influencing Ca_V_3.2 channel properties by chelation of trace metals such as zinc and copper ^26^ at the critical histidine residue. To explain the T-current facilitating properties of heat-treated and dialyzed serum, we searched for other endogenous Ca_V_3.2 channel modulators. Although the observed heat stability of the novel endogenous Ca_V_3.2 channel modulator discovered in this study might not be physiologically relevant by itself, this property indirectly led us to discover the low-MW peptide albumin (1-26) from the N-terminal end of albumin. Indeed, the N-terminus of albumin is an active site of this protein that binds and transports endogenous trace metals such as zinc ^32^.

Interestingly, albumin (1-26) and related peptides were previously found in the human urinary peptidome ^33^, and the human cerebrospinal fluid peptidome contains a shorter albumin fragment with two histidine residues within the first 10 N-terminal amino acids: albumin (1-10) ^34^. No previous reports, however, indicate that these peptides may act as nociceptive ion channel modulators. The present findings warrant further investigation to establish whether these and other albumin-related low-MW peptides in the human peptidome are implicated in pain disorders linked to neuronal hyperexcitability. This is particularly relevant given our findings that both native and recombinant human T-currents are effectively modulated by serum.

The existence of peptides like albumin (1-26) and/or other metal-chelating substances in the extracellular milieu of peripheral nociceptive DRG and DH neurons under physiological conditions is currently unknown. Previous studies reported that the endothelial cells vascularizing the DRG are unique, featuring larger fenestrations compared with those in proximal nerves, and are highly permeable to a wide range of low- and high-MW agents ^18^. Vascular permeability of the skin capillaries is important for tissue homeostasis and may be altered in various pathological conditions that allow leakage of various serum compounds ^35^. Hence, it is possible that inflammation, burns, tissue hematoma, surgical incisions, or other traumatizing events may result in serum extravasation and local accumulation of albumin itself or its N-terminal peptides (1-26) in the proximity of nociceptive neurons in DRGs and DHs, as well as in peripheral nerve endings. Prevailing literature suggests that trauma and disturbances in vascular integrity can induce pain by activating various inflammatory and metabolic mediators, such as cytokines, in the affected tissues ^36^. In contrast, we posit that direct modulation of a nociceptive ion channel by albumin itself or low-MW albumin fragments may sensitize both central and peripheral pain responses. Therefore, pharmacological targeting the mechanisms of endogenous modulation of Ca_V_3.2 channels and reversing neuronal hyperexcitability could offer therapeutic benefits for treating various pain disorders in humans. Importantly, we found that diluted human serum also potently modulated native T-currents in rat and human DRG neurons, and that T-currents in HEK-293 cells expressing the human variant of Ca_V_3.2 channels are also potentiated by serum, albumin, and the N-terminal albumin fragments. Therefore, we propose that the mechanisms of direct sensitization of pain responses by the endogenous modulators described herein may be highly relevant to human pain disorders.

Specifically, we found that serum augments the Ca_V_3.2 current density at very low concentrations over a wide range of test potentials, in addition to shifting the channel gating toward a more hyperpolarized potential. Both effects may promote T-channel activity by increasing its responsiveness to small depolarizations of the membrane and by facilitating the influx of calcium, which in turn decreases the threshold for I_Na+_ activation as demonstrated in the present study. To determine whether these *in vitro* electrophysiological effects had *in vivo* correlates, we investigated sensitivity to noxious heat and mechanical stimuli in rats and mice before and after i.pl. and i.t. injection of serum, albumin, and albumin (1-26). Blocking T-channels pharmacologically or using Ca_V_3.2 KO and H191Q Ca_V_3.2 KI mice eliminated the evoked hypersensitivity induced by the newly described modulators *in vivo*. These data confirm the results of our biophysical studies using DRG neurons *in vitro* and spinal cord preparations *ex vivo* and indicate that the serum modulators have a pro-nociceptive effect resembling inflammatory and/or neuropathic pain states that is mediated, at least in part, by direct modulation of Ca_V_3.2 channel function in peripheral and central pain pathways, respectively.

Our study provides definitive proof of the pro-nociceptive role of T-channel modulators by identifying their molecular structure and isolating them from the serum. Given the large number of different compounds present in serum, however, which is estimated to be >4000 ^37^, identifying a single molecule of interest is difficult. To begin, it was necessary to build a simple molecular profile by determining some of its key biochemical features using a variety of experimental methodologies. For example, because a protein’s structure governs its functional and chemical properties, we first characterized the mechanism by which it influences Ca_V_3.2 ion channel properties. Based on size exclusion dialysis, heat stability, stability in acidic conditions, as well as known sensitivity of Ca_V_3.2 channels to metal chelators by residue H191, we proposed and confirmed that the active serum component is a relatively low-MW, histidine-rich peptide that can act as an endogenous chelator of trace metals and a powerful modulator of neuronal excitability in pain pathways. Proteomic analysis, however, does not reveal all the proteins and peptides in a sample. Because of the sheer number of compounds, the mass spectrometry results are always compared with a known database. Therefore, all new peptides found are either already known peptides or fragments of known proteins. Our proteomic studies identified at least one more candidate peptide with a single histidine residue, such as VLSAADKGNVKAAWGKVGGHAAEYGAEALERM (Supplementary Figure 6, Table 2), that could also be expected to modulate Ca_V_3.2. It is possible that other currently unknown endogenous modulators of Ca_V_3.2 channels co-exist in the serum and cerebrospinal fluid and work in concert with albumin (1-26) identified in this study.

In conclusion, we found that recombinant Ca_V_3.2 currents and native Ca_V_3.2 currents *in vitro* and *ex vivo*, as well as thermal and mechanical nociception *in vivo,* are strongly stimulated by an endogenous serum peptide. These findings serve as a first step toward the development of pharmacological strategies to achieve long-term regulation of T-channel function in pain pathways. Elucidating the mechanisms of channel regulation will promote the development of specific therapies to treat diseases associated with the hyperactivity of T-channels and cellular hyperexcitability of nociceptive DRG and DH cells to reduce the availability of active T-channels, such as by gene therapy, i.t. delivery of monoclonal antibodies, or blocking peptides ^38,39^. This discovery opens novel avenues for pursuing novel analgesics with relatively unique mechanisms of action. We hope that this approach may greatly decrease the need for narcotics and the serious side effects associated with their use. Furthermore, the discovery of potent endogenous metal chelators may provide important insights into a variety of other physiological processes and pathological conditions, including those associated with neuronal injury, Alzheimer’s disease, Menke’s disease, and Wilson’s disease, in which increases in extracellular metals are documented ^40–44^. Finally, Ca_V_3.2 channel gain-of-function mutations are also implicated in hyperexcitable states associated with certain human seizure disorders ^45^. Thus, our discovery of endogenous chelators of trace metals and potent modulators of neuronal Ca_V_3.2 channels may have far-reaching importance across various fields of medicine and neuroscience.

## Materials and Methods

### In vitro preparation of human embryonic kidney-293 cells

Human embryonic kidney (HEK-293) cells were stably or transiently transfected to express human Ca_V_3.2 channels, as described previously ^11,26^. Cells were grown in Dulbecco’s modified Eagle’s medium and Ham’s F-12 Nutrient Mixture supplemented with 10% fetal bovine serum (FBS), 100 U/ml penicillin, and 0.1 mg/ml streptomycin (0.1 mg/ml), and incubated at 37℃ with 5% CO_2_. Media exchange was performed every 2 days, with regular splitting after every 4–7 days (at 85%–90% confluency).

Prior to recording, cells were split and plated onto poly-D-lysine-coated glass coverslips and allowed to adhere for 2–24 h in an incubator.

### In vitro preparation of acutely isolated rat DRG neurons

The properties of T-currents in putative nociceptive DRG cells (soma diameter <35 μm) freshly isolated from adolescent rats of both sexes were examined according to previously described procedures ^7,21,46^. For patch-clamp recordings, cells were plated onto uncoated glass coverslips and placed in a culture dish.

### Ex vivo preparation of acute rat spinal cord slices

Male (9 to 12-week-old) Sprague Dawley rats (Charles River Laboratories) were pair-housed on a 12-h light/dark cycle with *ad libitum* access to food and water. Animals were cared for in accordance with guidelines of the Canadian Council for Animal Care, Carleton University, and the University of Ottawa Heart Institute.

The rats were deeply anesthetized with an intraperitoneal injection of 1.5–3 g/kg urethane (Sigma). Spinal cords were then quickly dissected by ventral corpectomy and immersed in ice-cold oxygenated protective sucrose solution (cutting solution, [in mM]: 50 sucrose, 92 NaCl, 15 D-glucose, 26 NaHCO3, 5 KCl, 1.25 NaH2PO4, 0.5 CaCl2, 7 MgSO4, 1 kynurenic acid, bubbled with 5% CO_2_/95% O_2_). Dorsal and ventral roots were removed and the lumbar region (L3–L6) was isolated under a dissection microscope.

The lumbar spinal cords were sliced through the dorsal horn into 300-μm thick sections in the parasagittal plane using a Leica VT1200S vibratome at an amplitude of 2.75 mm and a speed of 0.1-0.2 mm/s. Slices were then incubated in kynurenic acid-free cutting solution at 34°C for 40 min. Following kynurenic acid washout, the spinal cord slices were removed from the heated water bath and allowed to passively cool to room temperature.

### Electrophysiology

For recordings in HEK-293 and DRG cells, electrodes were pulled from borosilicate microcapillary tubes to a final resistance of 2–5 MΩ. Spinal cord neurons were patched using fire-polished borosilicate glass patch-clamp pipettes with a resistance of 5–11 MΩ.

To isolate T-currents, electrodes were filled with internal solution comprising 135 mM tetramethylammonium (TMA) hydroxide, 10 mM ethylene glycol tetraacetic acid (EGTA), 2 mM MgCl_2_, and 40 mM N-2-hydroxyethylpiperazine-N’-2-ethanesulfonic acid (HEPES), and titrated in a plastic dish to a pH of 7.15–7.25 using hydrofluoric acid. The HEK-293 culture medium was exchanged with external solution comprising 152 mM tetraethylammonium (TEA) chloride, 10 mM HEPES, and 2 mM CaCl_2_, and titrated to a pH of 7.4 with TEA hydroxide before rinsing repeatedly to ensure complete removal of the FBS present in the culture media. For recordings of isolated T-currents in DRG neurons, cells were perfused with an external solution comprising 152 mM TEA chloride, 10 mM HEPES, and 10 mM BaCl_2_, and titrated to a pH of 7.42 with TMA hydroxide.

The internal solution for recordings of high-voltage-activated (HVA) calcium currents and excitability testing in DRG cells contained 110 mM Cs-MeSO_4_, 14 mM creatine phosphate, 10 mM HEPES, 9 mM EGTA, 5 mM Mg-ATP, and 0.3 mM Tris-GTP, adjusted to pH 7.3 with CsOH. In the recording chamber, we used Tyrode’s solution containing 140 mM NaCl, 4 mM KCl, 2 mM MgCl_2_, 2 mM CaCl_2_, 10 mM glucose, and 10 mM HEPES, adjusted to pH 7.4 with NaOH. All recordings from DRG neurons were performed within 6 h of dissociation.

For recordings in spinal cord slices, the internal patch pipette solution contained 140 mM K- gluconate, 4 mM NaCl, 10 mM HEPES, 1 mM EGTA, 0.5 mM MgCl2·6H20, 4 mM Mg-ATP, and 0.5 mM Na2-GTP, with a pH of 7.2 and an osmolarity of 290 mOsm. The extracellular recording solution contained 125 mM NaCl, 20 mM D-glucose, 26 mM NaHCO3, 3 mM KCl, 1.25 mM NaH2PO4, 2 mM CaCl2, and 1 mM MgCl2.

Gigaohm seals were reached prior to cell opening to obtain the whole-cell configuration. Membrane resistance, access resistance, and cell capacitance were monitored during the formation of the patch and the subsequent recordings.

Cells were perfused with various pre-treated FBS and albumin solutions using multiple independently controlled gravity-driven plastic and glass syringes. To test the synthesized low- molecular weight (MW) peptides in stably transfected HEK-293 cells, we recorded baseline currents in a recording chamber filled with 294 μl external bath solution prior to adding 6 μl peptide stock solution at a concentration 50 times higher than the desired final concentration. (This method was selected due to the limited peptide stock available for recordings). All treatment solutions were prepared from stock solutions and diluted to the appropriate concentrations before experimentation.

To obtain current-voltage (IV) curves, cells were maintained at a holding potential (V_h_) of -90 mV and subjected to multiple depolarizing steps (V_t_) ranging from -70 mV to +25 mV. The duration of the voltage step was 320 ms. To obtain steady-state inactivation curves, cells were held at V_h_ = -90 mV, hyperpolarized or depolarized (-110 mV to -40 mV) for a duration of 3.5 s, and then brought to a V_t_ = -30 mV.

For the spinal cord lamina I electrophysiological recordings, lamina I neurons were identified under brightfield optics as those situated dorsal to the substantia gelatinosa and within two cell widths ventral to superficial white matter tracts.

Whole-cell patch recordings were established at −60 mV. Neurons were required to have an access resistance below 35 MΩ, and leakage currents less than −80 pA. Spontaneous excitatory postsynaptic currents (sEPSCs) were recorded for a minimum of 20 min using a Clampex 10.7 (Molecular Devices). The sEPSC events were detected using MiniAnalysis 6.0.7 (Synaptosoft Inc.). Root mean square noise was measured in a region with no sEPSCs. The detection factor was set at 5 times the root mean square noise level. Negative-going peaks were auto-detected and manually verified. Events were then grouped by minute for analysis. Following a control baseline period, 1% FBS (Sigma) was added to the external recording solution during recording and allowed to perfuse onto the slice. A group of slices was pretreated with 1 µM TTA-P2 (Alomone Labs), a selective T-type voltage-gated calcium channel inhibitor. At 10 min before patching, 1µM TTA-P2 was also added to the external recording solution for this group.

All *in vitro* and *ex vivo* experiments were conducted at room temperature.

### Human DRG preparation and recordings

#### Tissue donation

Human DRG were sourced from two anonymized organ donors (♂52 y/o, ♀68 y/o), confirmed neurologically deceased after a stroke, in compliance with protocols sanctioned by the French organ transplantation authority, Agence de la Biomédecine (DC-2014-2420). Body temperature was lowered with ice, and blood circulation was maintained for 3 hours before vertebral bloc removal. After organ removal for transplantation purpose, a spinal segment from thoracic level (T9) to the caudal end was removed in 1 piece, and DRGs were immediately dissected and maintained in ice-cold, oxygenated calcium/magnesium free HBSS/Glucose/pen-strep for less than 30 minutes until returning to the lab. Then 3 DRGs per donor were cleaned and minced in small pieces and incubated at 37°C in 8ml of HBSS/Glucose/pen-strep solution containing 4 mg/ml collagenase (Sigma), 10mg/ml dispase (Sigma) and 5 mM calcium under gentle agitation. Tissue digestion was estimated by looking at isolated DRG somas release after triturating one piece of tissue using a fire polished Pasteur glass pipet with a large aperture. If needed and additional 30 to 60 min incubation of the tissues was performed. Digestion was stopped by washing the tissues two times with HBSS without calcium, and to times with culture medium (Neurobasal A / B27 / L-glutamine / pen-strep). Trituration was performed with Pasteur pipettes of decreased tips diameters. Debris were allowed to settle down the tube for 2 minutes and cell suspension was collected, passed on a 100 µm cell strainer, and centrifuged at 500g for 5 minutes. Cells were suspended in culture media and seeded on plastic petri dishes coated with poly D-Lysin and laminin A and kept at 37°C in a humidified culture incubator. The culture media was not supplemented with any growth factor.

#### Electrophysiology

Patch clamp recordings were performed 6 hours to 2 days after plating of DRG neurons using an Axopatch 200B amplifier (Molecular Devices, Sunnyvale CA) and fire-polished borosilicate glass pipettes with a typical resistance of 1.5–2.5 MOhm. Macroscopic calcium currents were recorded at room temperature using an internal solution containing (in mM): 100 CsCl, 40 tetraethylammonium (TEA)-Cl, 10 EGTA, 10 HEPES, 3 Mg-ATP, 0.6 GTPNa, and 3 CaCl2 (pH adjusted to 7.25 with KOH, ∼300 mOsm, ∼100 nM free Ca2+ calculated with the MaxChelator software) and an extracellular solution containing (in mM): 140 TEA-Cl, 10 4-Aminopyridin, 5 KCl, 2 CaCl2, 2 NaCl, 1 MgCl2, 10 HEPES and 10 Glucose (pH adjusted to 7.25 with TEA-OH, ∼310 mOsm). Extracellular solutions were applied by a gravity-driven homemade perfusion. Recordings were filtered at 5 kHz. Data were analyzed using pCLAMP10 (Molecular Devices) and GraphPad Prism 9 (GraphPad) software. Results are presented as the mean ± SEM, and n is the number of cells whereas N is the number of organ donors.

#### Chemical reagents

Compounds were purchased from Sigma (France). The effect of human serum (Sigma, H4522) was tested on human DRGs.

### Generation of H191Q Ca_V_3.2 knock-in (KI) mouse line

Guide RNA design was performed using CRISPOR and the Broad Institute sgRNA Design software, both of which have been refined to better identify off-target events and ensure more accurate gene editing ^47,48^. Results using each software were compared, and guides performing best using both algorithms were selected. Guide activity was verified by incubating guide RNA and Cas9 protein with a polymerase chain reaction (PCR) product containing the target sequence and comparing the ratio of cut to uncut PCR product. C57Bl/6J zygotes were electroporated with guide RNA (target sequence ATGATGGAATACTCTCTGGA CGG), Cas9 protein (Alt-R® S.p. HiFi Cas9 Nuclease V3 cat # 1081060), and a single-stranded DNA oligonucleotide (GAGACAGAGGCCTCTGGGCTCTTCCCTCCCTGAGCATCCCTGCTTCCCCAGCATGATGGA ATAtTCTCTaGACGGACAgAACGTGAGCCTCTCTGCCATCCGAACAGTGCGTGTGCTGCGGCCCCTCCGCGCCATCAACCG, IDT). Zygotes were then transferred into pseudopregnant recipients, resulting in the birth of 21 F0 pups from a single day’s procedure. The pups were genotyped by PCR using primers flanking the targeted region, followed by restriction enzyme digestion to identify putative positive founders. Of the 21 F0 founders born, 6 putative positive mice were identified. F0 founders were bred to B6J wild-type (WT) females and F1 offspring were sequence-verified to confirm the desired mutation. Homozygous mice from line 10 were transferred to the investigator for analysis.

### Behavioral studies *in vivo*

For behavioral studies, we used adult female rats and mice of both sexes. Rodents were maintained on a 14-h light/10-h dark cycle with free access to food and water. Experiments were conducted in accordance with institutional guidelines of the University of Colorado Anschutz Medical Campus and federal guidelines, including the National Institutes of Health *Guide for the Care and Use of Laboratory Animals* (NIH Publication No. 8023, revised 2002).

### Local intraplantar (i.pl.) injections in rats and mice

To test the behavioral effects of serum in rats, solutions containing 1% and 3% serum or vehicle in 100 μL of saline were intradermally injected into the ventral side of one hind paw. Intact albumin and albumin (1-26) were tested in rats in the same way using a concentration of 200 μmol/L. To test the behavioral effects of serum in mice, solutions containing 1% serum or vehicle in 20 μL of saline were intradermally injected into the ventral side of the right hind paw. All solutions were pH balanced to 7.4 to avoid skin irritation. No signs of skin inflammation, discoloration, or irritation were noted at the injection sites.

### Intrathecal (i.t.) injections in mice

To study the antinociceptive effects of drugs applied to the spinal cord, i.t. injections of the study compound were administered. After anesthetizing animals with isoflurane (2%–3%), the back of each animal was shaved to expose the injection site in the L4-L6 region of the spinal column.

After inserting the needle (28–30 G) into the L4-L6 region, 10 μl of the experimental compound or vehicle was delivered i.t. and the animal was allowed to recover for 20 min before initiating the experiments.

### Thermal paw withdrawal latency assay in rats

To assess the thermal nociceptive response to intraplantar injections of the test solution (100 μL), thermal paw withdrawal latencies (PWLs) in response to noxious heat stimuli were measured in rats using a custom-built paw thermal stimulator based on Hargreaves’s test. A moveable light source was positioned underneath either the right or left hind paw, as described previously ^24,25,49^. Timing began as soon as the light was turned on and ended as soon as the rat withdrew its paw. In all experiments with i.pl. injections, the uninjected paw served as a control.

### Mechanical hypersensitivity assay in mice

To test the mechanical sensitivity of the animals, we used the electronic Von Frey apparatus (Ugo Basile, Varese, Italy), which consists of one single rigid probe that exerts pressure in a range of 0–50 g. Animals were placed in plastic enclosures on a wire mesh stand to habituate for 15 min. After habituation, a probe was applied to the plantar surface of the paw through the mesh floor of the stand, and constant force was applied to the mid-plantar area of the paw. As soon as a rapid paw withdrawal occurred in response to the punctate stimulus, the apparatus displayed the force in grams, indicating the paw sensitivity threshold. Any other voluntary movement of the animal was not considered a response. In all experiments, 20 μl of test solution (serum or vehicle) was injected into one paw of gently restrained mice, while the uninjected paw served as a control. For i.t. injections, we injected 10 μl of test solution (serum or vehicle) into the lumbar spinal region under isoflurane anesthesia. The investigator assessing the behavioral responses was blinded to the pharmacological interventions.

### Drugs, chemicals, and peptides

Undialyzed and dialyzed FBS were purchased from Sigma-Aldrich and Dundee Cell Products, respectively. Human serum and horse serum were obtained from Sigma-Aldrich. Human cerebrospinal fluid was acquired from Lee Biosolutions. The T-channel blocker 3*β*-OH [(3*β*,5*β*,17*β*)-3-hydroxyandrostane-17-carbonitrile] was synthesized as described elsewhere ^19^. TTA-P2 (3,5-dichloro-*N*-[1-(2,2-dimethyl-tetrahydro-pyran-4-ylmethyl)-4-fluoro-piperidin-4- ylmethyl]-benzamide) was purchased from Alomone Labs. Using their amino acid sequence as a template, the active peptides were synthesized by automated solid-phase methods as described below.

### Serum purification and proteomics analysis

#### Serum preparation

Undialyzed FBS was treated with chloroform and heat. The chloroform treatment was a 1:1 solution of chloroform and FBS, which was vortexed and incubated on ice at 2-8°C for 24 h. The FBS was then centrifuged three times for 2 min each, and the separated aqueous phase was collected with a pipette. The FBS was left at 2-8°C for another 24 h to allow any remaining chloroform to evaporate. Heat treatment involved exposure to 100°C for approximately 15 min in a water bath. In a separate set of experiments, the aqueous FBS was subjected to dialysis using filtration columns with different MW cut-offs (Amicon Ultra-15 Centrifugal Filters, Sigma-Aldrich, MO) following the manufacturer’s instructions. Centrifugation was used to obtain FBS fractions with different MWs.

### Sample clean up

The peptides were desalted using C18 stop-and-go-extraction tips (StageTips). StageTips were prepared by punching 3M Empore C18 material with a Hamilton 20-gauge needle and inserting the disk into P200 pipette tips. StageTips were conditioned with 200 µL of 80% acetonitrile in 0.1% formic acid and then centrifuged for 1 min at 1500 x g. The StageTips were then washed with 200 µL 0.1% formic acid and transferred to clean microfuge tubes. Formic acid (0.1%) was then added to the concentrated sample to a final volume of 200 µL before vortexing for 15–20 min. The samples were added to the equilibrated C18 columns and centrifuged for 1 min at 1500 x g. The flow-through was reapplied to the resin three additional times to maximize sample binding. The StageTips were washed twice with 200 µL of 0.1% formic acid and centrifuged at 1500 x g for 1 min. The StageTips were transferred to a clean microcentrifuge tube and subjected to two rounds of elution with 100 µL of 0.1% formic acid in 80% acetonitrile, followed by centrifugation at 1500 x g. The eluent volume was reduced to near dryness using a Speedvac, and then resuspended with 10 µL of 0.1% formic acid for liquid chromatography-mass spectrometry analysis.

### Liquid chromatography-mass spectrometry analysis

Samples were analyzed on a Q Exactive quadrupole orbitrap mass spectrometer (Thermo Fisher Scientific, Waltham, MA, USA) coupled to an Easy nLC 1000 UHPLC (Thermo Fisher Scientific) through a nanoelectrospray ion source. Peptides were separated on a custom-made 20-cm C18 analytical column (100 µm x 10 cm) packed with 2.7 µm Cortecs C18 resin. After equilibration with 3 µL of 5% acetonitrile and 0.1% formic acid, the peptides were separated using a 120-min linear gradient from 4% to 30% acetonitrile with 0.1% formic acid at 400 nL/min. The liquid chromatography mobile phase solvents and sample dilutions used 0.1% formic acid in water (Buffer A) and 0.1% formic acid in acetonitrile (Buffer B) (Chromasolv LC–MS grade; Sigma- Aldrich, St. Louis, MO). Data acquisition was performed using the instrument’s Xcalibur™ (version 3.0) software. The mass spectrometer was operated in the positive ion mode using data- dependent acquisition. Full mass spectrometry scans were obtained with an m/z range of 300 to 1800, a mass resolution of 60,000 at m/z 200, and a target value of 1.00E+06 with a maximum injection time of 50 ms. High-energy collision dissociation was performed on the 15 most significant peaks, and tandem mass spectra were acquired at a mass resolution of 15,000 at m/z 200 and a target value of 1.00E+05 with a maximum injection time of 100 ms. Precursor ions were isolated using a window of 2 Th, meaning the instrument selected ions within 2 mass units around the target ion. The dynamic exclusion time was set to 20 s, and the normalized collision energy was 30.

### Peptide synthesis, purification, and high-resolution mass spectrometry

All standard solvents and reagents were obtained from Sigma-Aldrich unless otherwise specified. Fmoc-protected amino acids were purchased from New England Peptide, H-Rink-ChemMatrix resin (loading level: 0.47mmoml/g) from Gyros Technologies, trifluoroacetic acid from EMD Millipore, piperidine from Alfa-Aesar, 1-hydroxy-6-chloro-benzotriazole (6-Cl-HOBt) from Creosalus-Advanced Chemtech, and tri-isopropylsilane from Tokyo Chemical Industry (TCI). Analytical chromatography was performed on an Agilent 1100 model system equipped with an autosampler using Phenomenex columns as specified below. Preparative chromatography was conducted on a Waters Model 2525 binary pump system equipped with a Model 2487 absorbance detector. Mass spectral data were collected on an Orbitrap Fusion instrument.

All peptides were assembled by automated solid-phase methods using an Applied Biosystems 433A synthesizer. A standard *α*-fluorenyl-methoxycarbonyl/t-butyl (Fmoc/tBu)-based protecting group scheme was employed, using the following side chain protecting groups: Arg(Pbf), Asn(Trt), Asp(OtBu), Cys(Trt), Glu(OtBu), Gln(Trt), His(Trt), Lys(Boc), Ser(tBu), Thr(tBu), Tyr(tBu). The first residue was incorporated by coupling Fmoc-Asp-α-OtBu-ester via its *β*-carboxyl side chain to the Rink-ChemMatrix support. The initial and subsequent coupling cycles employed 6-Cl- hydroxybenztriazole (6-Cl-HOBt)/ diisopropylcarbodiimide for activation, with 20% piperidine in dimethylformamide for deprotection. Resin cleavage and side-chain deprotection were performed in 15 ml of trifluoroacetic acid containing 2% (v/v) of the following scavengers: water, thioanisole, 2-mercaptoethanol, and tri-isopropylsilane, for 2 h. The crude peptide was recovered by adding excess cold diethyl ether, followed by several washes of the precipitate with diethyl ether and air drying.

Purified peptide was isolated by preparative high-performance liquid chromatography using a Phenomenex Luna® 10 μm C8(2) 250 x 21.2 mm LC column and a 15%–30%B gradient over 45 min. The homogeneity of the purified peptides was assessed by analytical high-performance liquid chromatography using a Phenomenex Gemini® 5μm C18 150 x 4.6 mm analytical column and a linear gradient of 10%-40% B over 30 min.

For high-resolution mass spectrometry analysis, samples were suspended in 10 μL of 5% (v/v) acetonitrile, 0.1% (v/v) trifluoroacetic acid, and then directly injected onto a Waters M-class Acquity BEH C18 column (130 Å, 1.7 μm, 0.075 mm × 250 mm) at a flow rate of 0.3 μL min−1 using a Thermo Fisher Scientific nLC1000 system. Mobile phases used were (A) 0.1% formic acid aqueous and (B) 0.1% formic acid acetonitrile. Peptides were gradient eluted from 3% to 20% B in 24 min, 20% to 32% B in 5 min, and 32% to 85% B in 1 min. Mass spectrometry analysis was performed using an Orbitrap Fusion (Thermo Fisher Scientific). Samples were run in positive ion mode with a source spray of 1800 V and an ion transfer tube temperature of 275°C. MS1 scans were performed in the Orbitrap mass analyzer set to a mass range of 350–2000 m/z at a resolution of 120,000 and a target automatic gain control of 4 × 105.

### Data analysis

The voltage dependence of steady-state activation was described with a single Boltzmann distribution: G(V) = Gmax/(1 + exp[-(V - V_50_)/k]), where Gmax is the maximal conductance, V_50_ is the half-maximal activation voltage, and k (units of millivolts) is the slope factor representing the steepness of the voltage dependence of the distribution. The voltage-dependence of steady-state inactivation was also fit to the Boltzmann equation: I(V) = Imax/(1 + exp[(V - V_50_)/k]), where Imax is the maximal current amplitude, V_50_ is the half-maximal inactivation voltage, and k (units of millivolts) represents the slope factor.

For excitability testing, we determined the membrane potential at which I_Na+_ was activated in DRG neurons using IV curves with 2-mV incremental steps, both before and after FBS application, as well as during co-application of FBS with a T-channel blocker 3*β*-OH or TTA-P2. In each condition, we performed three successive recordings and calculated the average activation potential for each DRG neuron. In the spinal cord neuron experiments, the declared group size represents the number of independent measurements, and statistical analysis was performed using these values.

Due to the considerable variability in sEPSC frequencies between cells, the data for each cell were normalized to the average of a 3-min baseline period. For saline-treated cells, this baseline corresponded to minutes 3, 4, and 5 of the recording. For drug-treated cells, the baseline was the 3 min preceding the drug application, which was typically administered at the 5-min mark of the recording. To evaluate the effect of FBS, 4-min averages of the normalized sEPSC frequency were calculated starting 8 min after the start of the treatment period. To account for the variable onset of effect, the average of the 4-min period with the highest frequency was selected for each treatment (saline, FBS, TTA-P2+FBS) and used for statistical analysis.

Comparison of means was conducted using IBM SPSS Statistics 28. No outliers were omitted in this dataset. Assumptions of normality (tested by Shapiro-Wilk) and homogeneity of variance (tested by Levene’s Test) were assessed. The nonparametric Kruskal-Wallis test was applied when the assumption of normality was not met, followed by pairwise comparisons with Bonferroni-adjusted significance when significance was reached in the Kruskal-Wallis test. After analysis, traces were transferred to Origin Pro (Northampton) for graphing.

Where not stated otherwise, statistical significance was determined using two-way analysis of variance (ANOVA). P-values <0.05 were considered significant. P-values and standard errors of the mean (SEM) from multiple determinations are provided in the text.

For all behavioral experiments, statistical comparisons were made using two-way repeated measures ANOVA, followed by the Holm-Sidak or the Bonferroni multiple comparisons, with statistical significance set at p < 0.05.

## Acknowledgments

We would like to thank Drs. Paula Barrett and Junlan Yao at the University of Virginia in Charlottesville for providing the stably transfected HEK-293 cells. We thank Dr. Kathiresan Krishnan for preparing 3*β*-OH in the laboratory of Dr. Douglas Covey at Washington University in Saint Louis. We thank Dr. Monika Dzieciatkowska, Mr. Mark Maslanka, and Dr. Kirk Hansen for their contributions through the Proteomics Core at the University of Colorado, Anschutz Medical Campus. We also thank Dr. Jennifer Matsuda and the Regional Mouse Genetics Core Facility at National Jewish Health, Denver, CO. We gratefully acknowledge the gift of human tissue from all donors included in this study and their families, whose contribution has been critical.

