## SupplementalFigures1-10 for "Facilitation of Ca_V_3.2 channel gating in pain pathways reveals a novel mechanism of serum-induced hyperalgesia"

**Supplemental Figure 1. Human serum potentiates T-currents in rat DRG neurons in a voltage--dependent manner.**

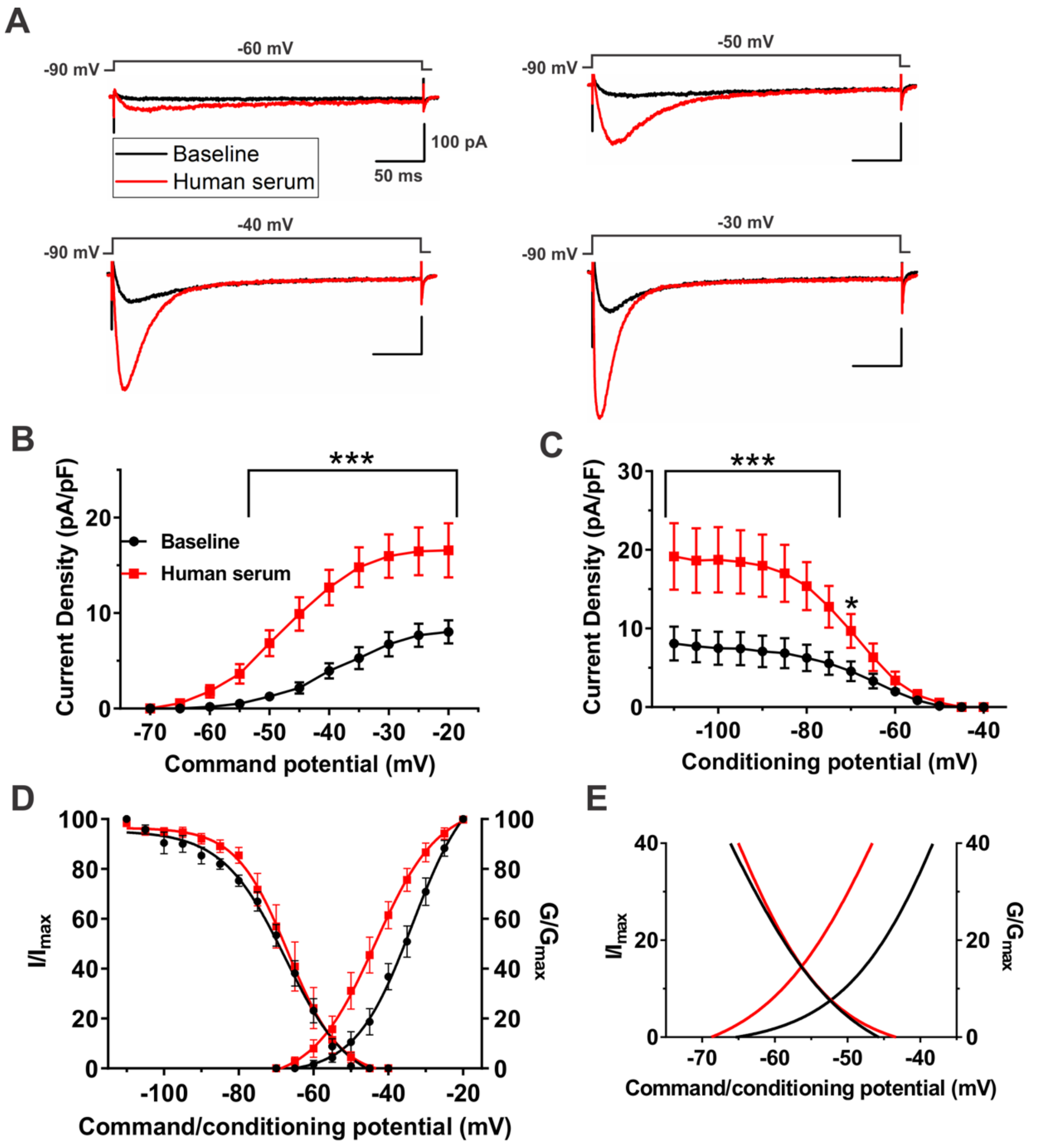


**(A)** Original traces of T-currents using IV protocols from a representative DRG neuron before (black) and after the addition of 1% human serum (red trace) in the voltage range of $V_t$ from -60 (top left) to -30 mV (bottom right) from $V_h$ of -90 mV (top).

**(B)** Voltage dependence of activation expressed in T-current densities before (black) and after the addition of 1% human serum (red). Two-way repeated measures ANOVA followed by Sidak's multiple comparisons test; interaction: $F(10,60)=11.49$, $p<0.001$; Sidak's: $p<0.001$ for potentials from -50 to -20 mV; n=7 neurons, 2 rats.

**(C)** Average current densities of the voltage dependence for steady-state inactivation before (black) and after the addition of 1% human serum (red). Interaction: $F(14,70)=8.69$, $p<0.001$; Sidak's: $p<0.001$ for potentials from -110 to -75 mV, and $p=0.032$ for -70 mV; n=6 neurons, 2 rats.

**(D)** Graph shows both normalized voltage dependence of activation ($G/G_{max}$) and steady-state inactivation ($I/I_{max}$) before (black) and after the addition of 1% human serum (red).

**(E)** The overlap of $G/G_{max}$ and $I/I_{max}$ curves shown in panel D yielded an area known as the window current. Note that this current, which allows for a steady influx of calcium near the resting membrane potential, was increased and shifted to the left upon the addition of human serum. Data are presented as mean ± SEM. *$p<0.05$ and ***$p<0.001$ compared to baseline.

**Supplemental Figure 2. Human Serum selectively increases Peak T-type current in acutely dissociated human DRG cells.**

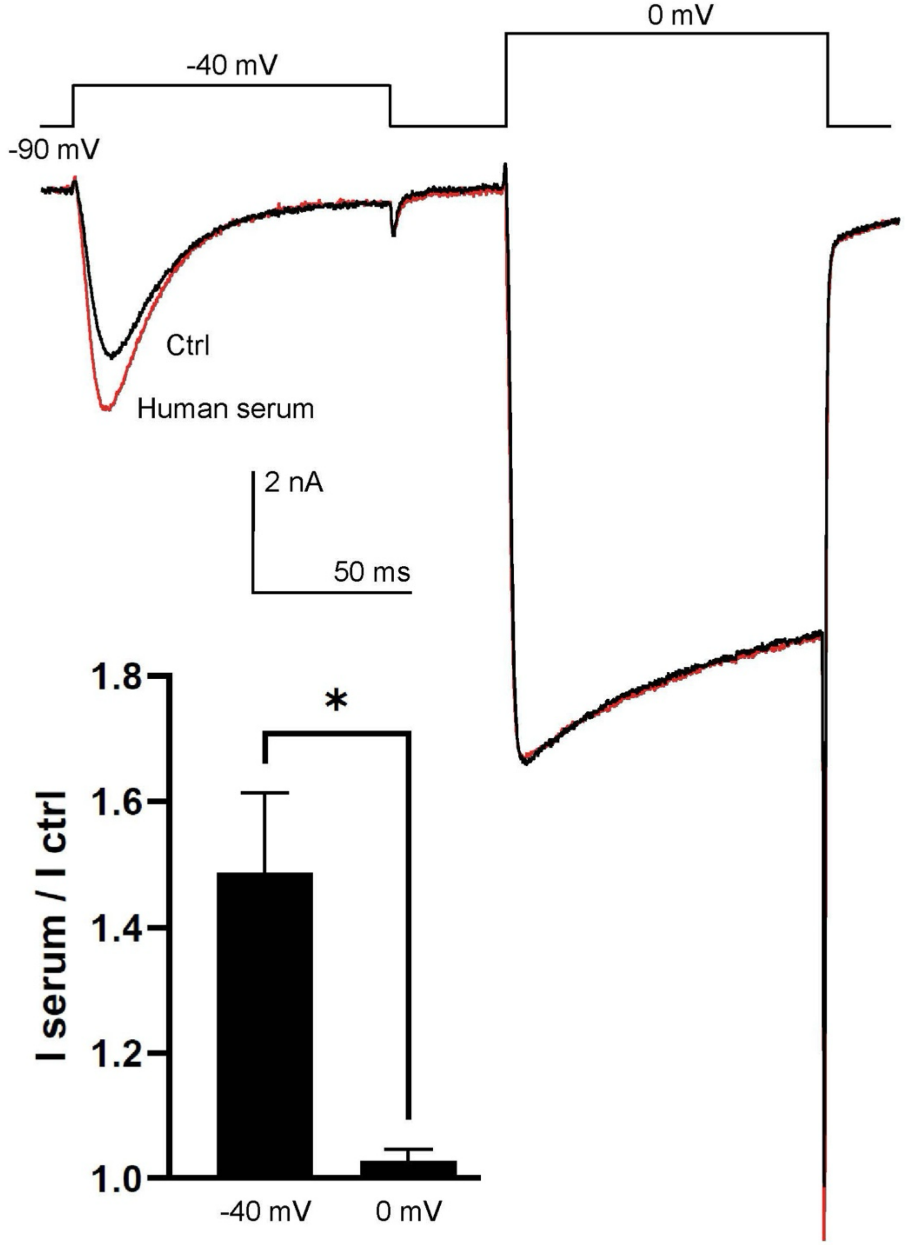


Representative traces of T-type current $V_h$ -90 mV, $V_t$ -40 mV and high-voltage-activated (HVA) calcium current ($V_h$ -90 mV, $V_t$ 0 mV) for an acutely dissociated human DRG cell. One percent human serum increased the T-type current recorded at -40 mV (I serum /I ctrl = 1.485 +- 0.1282 (mean+-SEM) but not the HVA current at 0 mV (0.1282 +- 0.01913). This effect of serum on T-type current is significant as compared to HVA current (p=0.0286, non-parametric Mann Whitney test). Error bars represent SEM. * = p<0.05 (n=4 cells from 2 donors).

**Supplemental Figure 3. Serum increases sEPSC frequency in rat dorsal horn (DH) neurons in the acute spinal cord slice preparation.**

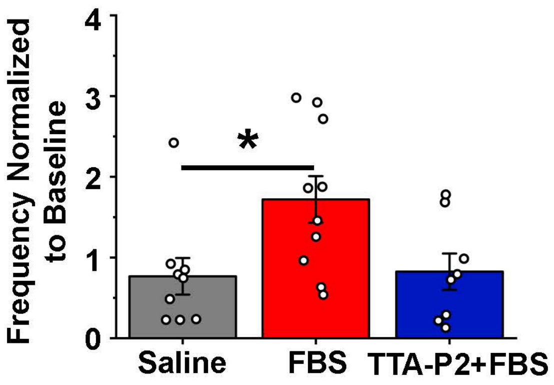


Cumulative frequency normalized to baseline (calculated by taking the highest 4-minute average period of normalized frequency for each cell 8-22 minutes following the start of treatment, thus assuring no bias in selecting a region, but accounting for the variability in the onset of effect of FBS on sEPSC frequency) for saline (n=9, gray), 1%FBS (n=10, red), and 1μM TTA-P2 (3,5-dichloro-N-[1-(2,2-dimethyl-tetrahydro-pyran-4-ylmethyl)-4-fluoro-piperidin-4-ylmethyl]-benzamide) + 1%FBS (n=8, blue). Error bars represent SEM. Kruskal-Wallis test followed by pairwise comparison with Bonferroni adjusted significance. * = $p<0.05$

**Supplemental Figure 4. Gene editing of cacna1h for generation of H191Q $Ca_V3.2$ KI mouse.**
Cacna1h genomic DNA organization and predicted isoforms are shown below (Ensembl). Transcript -201 was used to identify H191 within the coding sequence for guide RNA and HDR template design.

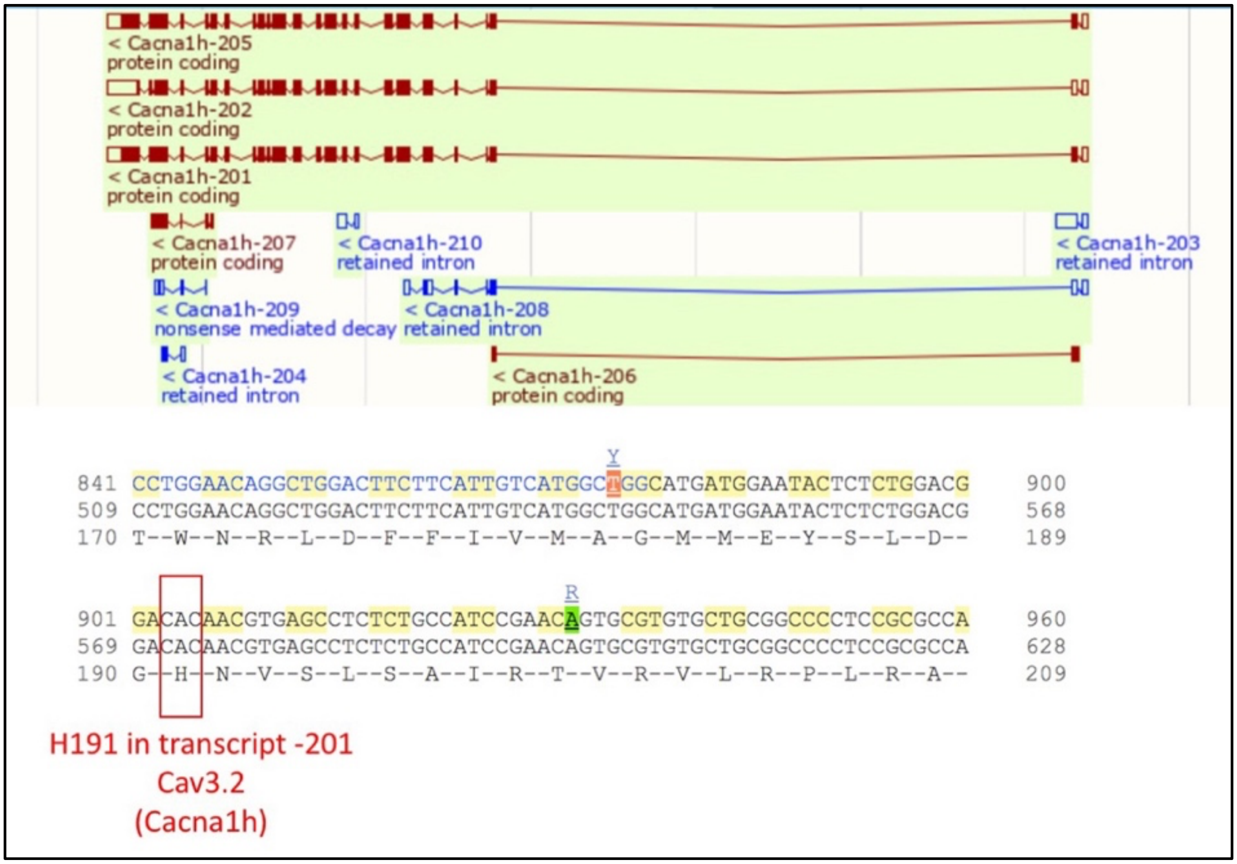

| | Number of cells *N* | $I_{max}$ Fold-Increase | SEM | *P*-value | Summary |
|---|---|---|---|---|---|
| FBS 1% | 36 | 1.92 | 0.07 | <0.0001 | **** |
| Heat-treated FBS 1% | 3 | 2.03 | 0.26 | <0.0001 | **** |
| FBS exposed to pH < 2 | 5 | 1.76 | 0.07 | <0.0001 | **** |
| FBS exposed to pH >10 | 5 | 1.07 | 0.05 | 0.9995 | ns |
| FBS + copper | 7 | 1.12 | 0.21 | 0.9576 | ns |
| Hydrophylic fraction of FBS | 4 | 2.40 | 0.24 | <0.0001 | **** |
| FBS fraction >1 kDa | 10 | 2.01 | 0.08 | <0.0001 | **** |

**Supplemental Figure 5**
**Table 1. Various steps of purification of serum indicate that the T-channel modulating substance is a heat-stable, acid-stable, and hydrophilic molecule.**

Copper was added to FBS at a concentration of 10 μM, FBS was exposed to treatment at a concentration of 100% and diluted 1:100 with the external patch clamp solution prior to perfusion. P-values were determined using a paired mixed-effect analysis with Dunnett's multiple comparison's test comparing each value to the baseline current recorded from the same putative nociceptive DRG neuron (treatment: $F(2.513,35.18) = 32.36$, $p<0.0001$).

**Supplemental Figure 6: Active fraction of fetal bovine serum (FBS) has approximate molecular weight (MW) of 3-5 kDA.**

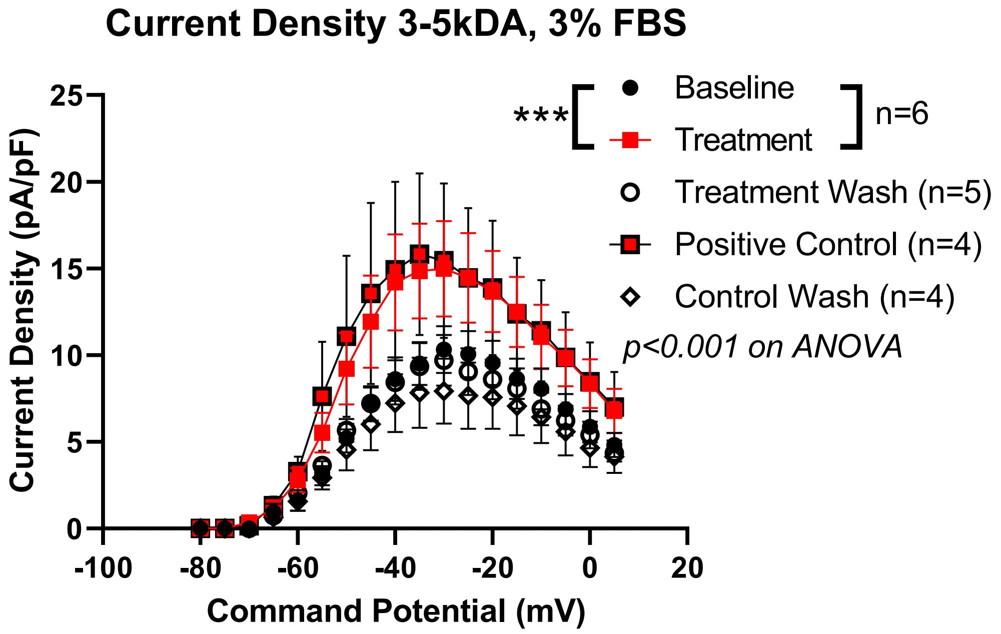


Averaged current-voltage (IV) curve recorded from $Ca_V3.2$ channel-expressing HEK-293 cells show that FBS (red symbols) increased the amplitudes of peak current densities when compared to the control pre-drug conditions (black symbols) over a wide range of potentials (V*h* = -90 mV, V*t* = -80 mV through 5 mV in 5 mV increments).
In each recording baseline $Ca_V3.2$ current amplitudes ($V_h$ -90 mV, $V_t$ -30 mV) were determined before application of either positive control with 3% FBS or treatment with 3% of solution dialyzed against 3-5 kDA membrane cut off in the same cells. Peak currents densities were obtained by normalizing peak current amplitudes (in pA) to cell capacitance (in pF). Notice than both “positive control” and “treatment” caused virtually identical and fully reversible increase in the $Ca_V3.2$ current densities over the wide range of potentials. Number of cells is indicated in parenthesis, symbols are averages of multiple determinations, vertical lines represent ± SEM.
*** p<0.001 on ANOVA

**Methods for preparation of 3-5 kDa serum fraction**:

1. <u>Create a 3kDa fraction</u>

i. Prepare 3kDa 15ml size filter (max starting volume 15ml)

1. Rinse to remove glycerin with MilliQ water, keep wet

ii. Load undialyzed sample into filter

1. Spin in swinging bucket centrifuge at 4000G at 22C for 60 min

iii. Remove TOP fraction of filter, insert into the following:

2. <u>Create a 3-5kDa fraction</u>

i. Prepare GE Healthcare Vivaspin 20 5kDa MWCO filter (5-20ml capacity)

1. Run 20 ml Di H20 through 1000G at room temp for 1 min
2. Run about 5ml ethanol 70% through 1000G for 1 min

ii. Load 3kDa sample into filter

1. Spin 4000G at room temp for 30 min

**Supplemental Figure 7:**

The 10 most abundant peptides revealed by MS proteomic analysis of the active fraction of partially purified fetal bovine serum (FBS).

| Peptide | Length | Mass (Da.) | AUC Rep 1 | AUC Rep 2 | AUC Rep 3 | average AUC |
|---|---|---|---|---|---|---|
| YDRNTKSPLFVGKVVNPTQA | 20 | 2233.19 | 1.87E+10 | 1.48E+10 | 1.59E+10 | 1.64E+10 |
| VLSAADKGNVKAAWGKVGGHAAEYGAEALERM | 32 | 3255.65 | 2.83E+09 | 2.56E+09 | 3.96E+09 | 3.12E+09 |
| AAIDEASKKLNAQ | 13 | 2989.58 | 1.42E+09 | 1.63E+09 | 7.71E+08 | 1.27E+09 |
| GESGREGAPGAEGSPGRDGSPGAKGDRGETGP | 32 | 1357.72 | 1.18E+09 | 3.64E+08 | 5.64E+08 | 7.03E+08 |
| DTHKSEIAHRFKDLGEEHFKGLVLIA | 26 | 2999.32 | 1.47E+09 | 2.73E+08 | 1.08E+08 | 6.18E+08 |
| GKNGDDGEAGKPGRPGERGPPGP | 23 | 2233.05 | 1.16E+09 | 2.01E+08 | 2.56E+08 | 5.40E+08 |
| SEETKENERFTV | 12 | 2048.91 | | 6.14E+08 | 6.26E+08 | 6.20E+08 |
| EDGSDPPSGDFLTEGGGV | 18 | 1250.55 | 1.51E+08 | 1.96E+08 | 8.52E+08 | 4.00E+08 |
| PPSGDFLTEGGGV | 13 | 1231.57 | 3.31E+08 | 4.40E+08 | 3.26E+08 | 3.65E+08 |
| ADGQPGAKGEPGDAGAKGDAGPPGP | 25 | 2204.99 | 3.06E+08 | 2.21E+08 | 2.48E+08 | 2.58E+08 |

**Table 2:** The first column on this table shows peptide structure as determined by MS/MS of dialyzed fetal bovine serum fraction. Highlighted in yellow is albumin (1-26) N-terminal peptide with 3 histidine residues as denoted with red fonts. The second and third columns show peptide length and calculated molecular mass in Da, respectively. Columns 4-6 show AUC for three replicates and last column on the right shows average values of AUC from 3 replicates.

**Supplemental Figure 8. Heat treatment of bovine serum albumin (BSA) abolishes its $Ca_V3.2$ T-current enhancing properties.**

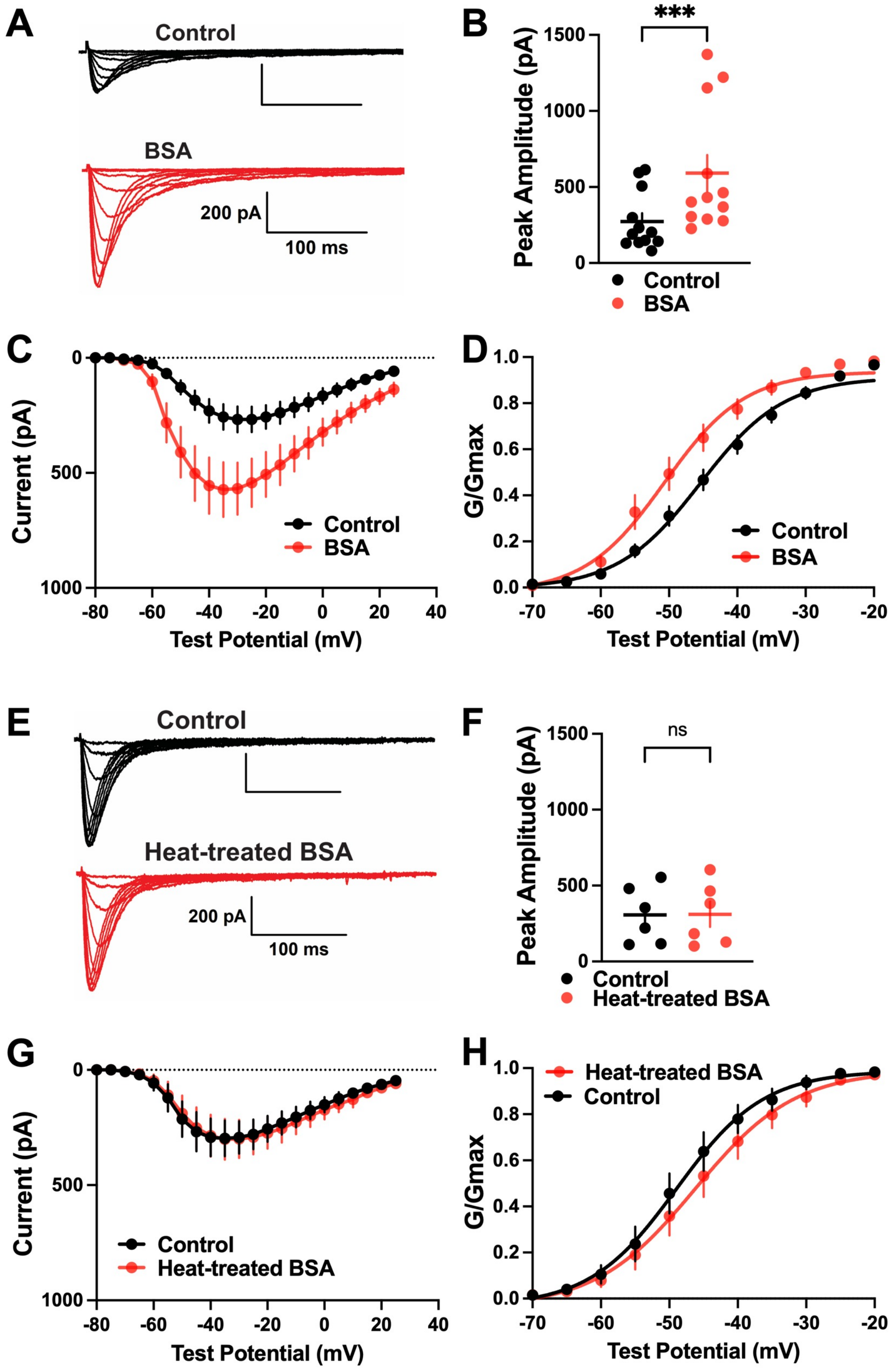

**(A)** Representative $Ca_V3.2$ T-current traces using IV protocols before (black) and after (red) the addition of 7 µM BSA to the bath solution recorded in stably transfected HEK-293 cells (V*h* = -90 mV, V*t* = -70 mV through -30 mV).

**(B)** Summary of the average maximal peak current amplitudes displayed as dot-plots (control mean amplitude: 273.2 ± 54.9 pA, BSA mean amplitude: 591.8 ± 118.4 pA, n=12, p<0.001, two-tailed paired student's t-test).

**(C)** The current-voltage relationship of $Ca_V3.2$ T-currents in control conditions (black) and after addition of BSA (red) in the same HEK-293 cells (n=12).

**(D)** Conductance-voltage relationship depicting the differences of channel activation as calculated using the Boltzmann function ($V_{50}$ control: -45.03 ± 1.33, $V_{50}$ BSA: -49.94 ± 1.63, p<0.001 two-tailed paired t-test).

**(E)** Representative recordings of $Ca_V3.2$ currents using the IV protocol before (black) and after (red) application of 7 µM heat-treated BSA (V*h* = -90 mV, V*t* = -70 mV through -30 mV).

**(F)** Average maximal peak current amplitudes in control conditions (black) and after application of 7 µM heat-treated BSA (red) in the same cells displayed as dot-plots (mean control: 306.5 ± 76.4 pA, mean heat-treated BSA: 311.0 ± 83.28, n=6, p=0.75, two-tailed paired t-test).

**(G)** The current-voltage relationship shows almost exactly overlapping curves before and after the application of heat-treated BSA.

**(H)** Comparison of activation kinetics using conductance normalized to maximal conductance fitted with the Boltzmann equation in control conditions (black) and after perfusion with 7 µM heat-treated BSA (red) ($V_{50}$ control: -48.46 ± 2.24 mV, $V_{50}$ heat-treated BSA: -45.71 ±2.55, n=6, p=0.002, two-tailed paired t-test).

**Supplemental Figure 9. The ability to augment $Ca_V3.2$ currents by N-terminal albumin peptide (1-26) is conserved between several mammalian species.**

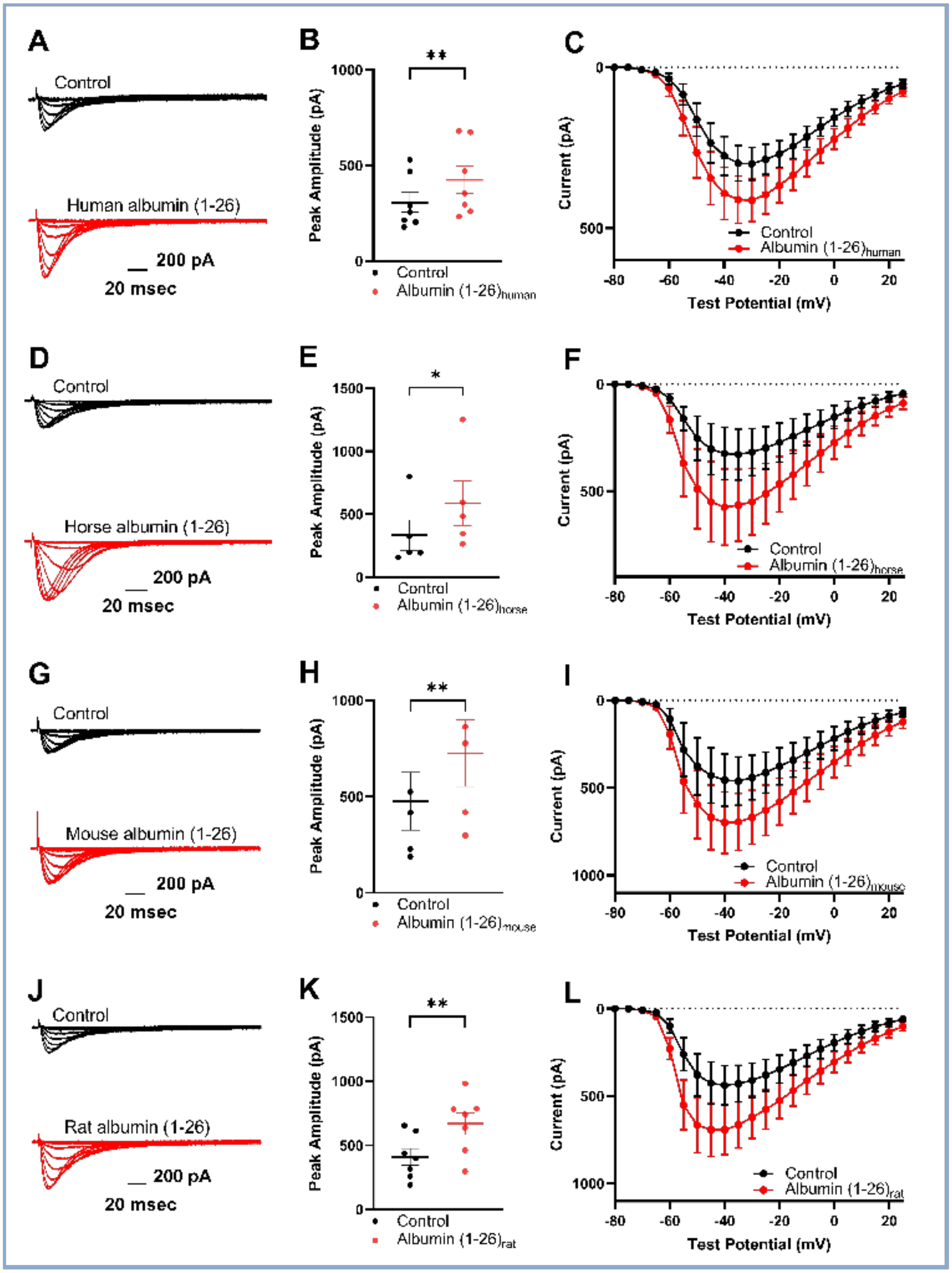

**(A)** Representative IV current traces before (black) and after application of 0.4 mM human albumin (1-26) peptide (red).

**(B)** The mean maximal peak current amplitude in control conditions was 305.7 ± 52.3 pA and increased after addition of human albumin (1-26) peptide to 423.9 ± 71.61 pA (n=7, p=0.002, two-tailed paired t-test).

**(C)** Current-voltage relationship before (black symbols and lines) and after addition of 0.4 mM human albumin (1-26) peptide (red symbols and lines).

**(D)** Representative IV current traces before (black) and after application of 0.4 mM horse albumin (1-26) peptide (red).

**(E)** The mean maximal peak current amplitude in control conditions was 333.8 ± 119.4 pA and increased after addition of equine albumin (1-26) peptide to 587.5 ± 175.6 pA (n=5, p=0.02, two-tailed paired t-test).

**(F)** Current-voltage relationship before (black symbols and lines) and after addition of 0.4 mM equine albumin (1-26) peptide (red symbols and lines).

**(G)** Representative IV current traces before (black) and after application of 0.4 mM synthetic mouse albumin (1-26) peptide (red).

**(H)** The mean maximal peak current amplitude in control conditions was 475.7 ± 149.8 pA and increased after addition of 0.4 mM mouse albumin (1-26) peptide to 725.8 ± 172.9 pA (n=5, p=0.006, two tailed paired t-test).

**(I)** Current-voltage relationship before (black symbols and lines) and after application of 0.4 mM mouse albumin (1-26) peptide (red symbols and lines).

**(J)** Representative IV current traces before (black) and after application of 0.4 mM rat albumin (1-26) peptide (red).

**(K)** The mean maximal peak current amplitude in control conditions was 410.1 ± 65.7 pA and increased after addition of 0.4 mM rat albumin (1-26) peptide to 669.7 ± 86.2 pA ($n=7$, $p=0.002$, two tailed paired t-test).

**(L)** Current-voltage relationship before (black symbols and lines) and after addition of 0.4 mM rat albumin (1-26) peptide (red symbols and lines).

**Supplemental Figure 10**. Corresponding amino acid sequences from human, horse, mouse, and rat albumin (1-26) peptides.

Table 1

| Peptide | Amino acid sequence |
|---|---|
| Albumin $(1\text{-}26)_{bovine}$ | NH2-DTHKSEIAHRFKDLGEEHFKGLVLIA-CONH2 |
| Albumin $(1\text{-}26)_{human}$ | NH2-DAHKSEVAHRFKDLGEENFKALVLIA-CONH2 |
| Albumin $(1\text{-}26)_{horse}$ | NH2-DTHKSEIAHRFNDLGEKHFKGLVLVA-CONH2 |
| Albumin $(1\text{-}26)_{mouse}$ | NH2-EAHKSEIAHRYNDLGEQHFKGLVLIA-CONH2 |
| Albumin $(1\text{-}26)_{rat}$ | NH2-EAHKSEIAHRFKDLGEQHFKGLVLIA-CONH2 |
